# Optimising 7T-fMRI for imaging regions of magnetic susceptibility

**DOI:** 10.1101/2025.03.17.643748

**Authors:** Saskia L. Frisby, Marta M. Correia, Minghao Zhang, Christopher T. Rodgers, Timothy T. Rogers, Matthew A. Lambon Ralph, Ajay D. Halai

## Abstract

The temporal signal-to-noise ratio (tSNR) of functional magnetic resonance imaging (fMRI) is particularly poor in ventral anterior temporal and orbitofrontal regions because of B_0_ and B_1_^+^ magnetic field inhomogeneity, a problem that is exacerbated at higher field strengths. In this 7T-fMRI study we compared three methods of improving sensitivity in these areas: parallel transmit, which uses multiple transmit elements, controlled independently, to homogenise the flip angle experienced by the tissue; multi-echo, which entails collection of multiple volumes at different echo times following a single radiofrequency pulse; and multiband, in which multiple slices are acquired simultaneously. We found that parallel transmit and multi-echo increased the magnitude of the BOLD signal change, but only multi-echo increased BOLD magnitude in areas prone to susceptibility artefacts. Multiband and denoising of multi-echo data with independent components analysis (ICA) both improved precision of GLM fit. Exploratory results suggested that multi-echo and ICA denoising can both benefit multivariate analyses. In conclusion, a multi-echo, multiband sequence improved fMRI quality in areas prone to susceptibility artefacts while maintaining sensitivity across the whole brain. We recommend this approach for studies investigating the functional roles of ventral temporal and orbitofrontal regions with 7T fMRI.

## 1. Introduction

The temporal signal-to-noise ratio (tSNR) of functional magnetic resonance imaging (fMRI) varies across the brain. The ventral anterior temporal cortex and orbitofrontal cortex, for example, are affected by two kinds of magnetic field inhomogeneity. First, these regions are located next to air-filled sinuses and so are affected by B_0_ inhomogeneity that causes signal dropout and geometric distortions (Devlin et al., 2000; Halai et al., 2014, 2015, 2024; Jezzard & Clare, 1999). The effects are more severe at higher field strengths, implying that it may be especially difficult to measure task-related activity in susceptible regions with ultra-high-field fMRI (e.g., 7T-fMRI). Second, at 7T and above, the radiofrequency wavelength is similar to the size of the head, resulting in standing wave effects that cause B_1_^+^ inhomogeneity in the temporal lobes (Gras et al., 2019; van der Kolk et al., 2013). This in turn causes spatial variation in flip angle and therefore in image intensity (Uğurbil, 2018; Wu et al., 2018). B_0_ and B_1_^+^ inhomogeneity make it challenging to use fMRI to investigate the roles that these regions may play in a myriad of cognitive processes – including vision (Devereux et al., 2018), language (Borghesani et al., 2016), multimodal semantic cognition (Lambon Ralph et al., 2017), emotion (Fernandez et al., 2017), social cognition (Binney et al., 2016; Zahn et al., 2007), theory of mind (DuPre et al., 2016), and executive function (Duncan, 2010). However, 7T-fMRI also has many advantages in regions unaffected by inhomogeneity: 7T-fMRI offers improved tSNR relative to 3T-fMRI (Morris et al., 2019), which can be used to reduce voxel size and enable applications such as laminar fMRI (Koopmans et al., 2011) or to reduce acquisition times and enable shorter scan times for special populations such as patients with neurodegenerative diseases (Cope et al., 2023). 7T-fMRI also benefits from improved spatial specificity relative to 3T-fMRI because the signal from cortical microvasculature is enhanced while the signal from large veins is reduced (Marques & Norris, 2018). Improving signal homogeneity would allow researchers studying the whole brain (or focusing on regions prone to susceptibility artefacts) to take full advantage of 7T-fMRI; therefore, in this study we compared three methods of doing so.

One possible method for improving image quality is parallel transmit (pTx), which uses multiple transmit elements, controlled independently, to homogenise the flip angle pattern experienced by the tissue (Adriany et al., 2005; Deniz et al., 2019; Roemer et al., 1990; Uğurbil, 2018; Van de Moortele et al., 2005). In this way, pTx directly counteracts the effect of B_1_^+^ inhomogeneity – however, since the design of spoke pTx pulses incorporates B_0_ field maps, pTx is also capable of counteracting B_0_ inhomogeneity to an extent (Zhang et al., 2024). A recent study used pTx to improve image quality in ventral anterior temporal regions for echo-planar imaging (EPI) 7T fMRI (Ding et al., 2022). pTx improved tSNR across the brain compared to a standard sequence, particularly in the temporal lobes. However, there was no improvement in functional contrast during a semantic association task that is known to recruit the anterior temporal lobes in 3T-fMRI studies (Jung et al., 2017).

A second method of recovering signal in these regions is multi-echo (ME) imaging (Kundu et al., 2017; Poser et al., 2006; Posse, 2012), which aims to counteract the effects of B_0_ inhomogeneity. T_2_* is known to vary across the brain (Hagberg et al., 2002); in areas affected by B_0_ inhomogeneity, T_2_* is particularly short due to increased intravoxel dephasing. A single echo provides sensitivity to a narrow range of T_2_* values; the echo time (TE) is therefore selected to provide the best compromise of sensitivity to T_2_* across the whole brain. Combining data from multiple echoes increases the range of T_2_* that can be imaged with high fidelity. ME has been shown to improve functional contrast (Poser & Norris, 2009) and spatial specificity (Boyacioğlu et al., 2015) at 7T and 3T (Fernandez et al., 2017; Halai et al., 2024; Kirilina et al., 2016; Lynch et al., 2020). Having multiple echoes also facilitates the separation of signal and noise because signals decay in a well-characterised way across echoes, whereas noise does not. This principle underpins multi-echo independent components analysis (ME-ICA), via which ICA components that are TE-independent, and thus are likely to be noise rather than blood-oxygen-level-dependent (BOLD) signal, can be removed (Dipasquale et al., 2017; Kundu et al., 2011, 2013, 2015, 2017). This method may enhance signal detection in areas prone to susceptibility artefacts on top of the advantage offered by ME alone (e.g. Lombardo et al., 2016). ME sequences have some potential disadvantages. For example, ME can lengthen repetition time (TR). In-plane acceleration is frequently needed to achieve a sufficiently short first TE, which reduces tSNR (Yun & Shah, 2017). In turn, a short first TE, combined with hardware constraints, often limits the minimum voxel size (Koopmans et al., 2011). Critically, in many previous studies examining the benefits of ME sequences compared to single-echo (SE), the “single-echo” data were extracted from the ME dataset (Amemiya et al., 2019; Bhavsar et al., 2014; Caballero-Gaudes et al., 2019; Cohen et al., 2017, 2017, 2018; Dipasquale et al., 2017; Evans et al., 2015; Fernandez et al., 2017; Gilmore et al., 2022; Kovářová et al., 2022). This means that the “single-echo” data will inherit suboptimal parameters that are ME-specific, making the comparison unfair.

A third strategy for improving acquisition is multiband (MB) imaging, also known as simultaneous multi-slice, in which multiple slices are acquired simultaneously (Barth et al., 2016; Moeller et al., 2010; Setsompop et al., 2012). Rather than counteracting B_0_ or B_1_^+^ inhomogeneity directly, MB has typically been applied to increase temporal resolution – multiple studies have shown that MB can reduce noise aliasing, increase statistical power and counteract the increase in TR associated with ME (Feinberg et al., 2010; Griffanti et al., 2014; Halai et al., 2024; Puckett et al., 2018; Smith et al., 2013). Benefits specific to task fMRI have been less clear (Demetriou et al., 2018; Todd et al., 2016) since some noise features are unlikely to be correlated with task regressors and can be removed with high-pass filtering. Other possible disadvantages of MB include a reduction in tSNR due to increases in g-factor effects (Demetriou et al., 2018; Risk et al., 2021; Setsompop et al., 2012) and leakage of signal into the simultaneously-excited slices (Todd et al., 2016).

Previous work therefore makes clear predictions about which 7T-fMRI sequences are capable of improving sensitivity in regions affected by B_0_ and B_1_^+^ inhomogeneity without compromising signal quality elsewhere. In this study we tested five possible sequences with the goal of demonstrating their *relative* impacts on functional effects of interest. These consisted of a 2 x 2 factorial design varying number of echoes and multiband factor – single-echo single band (SESB), single-echo multiband (SEMB), multi-echo single band (MESB), and multi-echo multiband (MEMB) – plus a fifth sequence that used pTx. Rather than attempting to map the limits of each parameter, our aim was to generate sequences that would be of practical benefit to neuroscientists seeking to answer a range of cognitive questions that implicate regions affected by inhomogeneity.

We used a semantic judgment task that is known to evoke activity across the semantic network, including areas severely affected and those relatively unaffected by susceptibility artefacts (Jung et al., 2017).

We had two univariate effects of interest – activation magnitude and activation precision (Halai et al., 2024). Activation magnitude is the magnitude of the BOLD signal change, operationalised as the 1^st^-level beta values extracted from each voxel. We hypothesised that both pTx and ME would recover signal and hence increase activation magnitude relative to the SESB sequence (which we used as a baseline). Activation precision is the reliability of the BOLD signal change, analogous to the functional contrast-to-noise ratio (fCNR) and operationalised as the 1^st^-level t-values extracted from each voxel. We hypothesised that MB sequences, with greater effective degrees of freedom, would increase activation precision relative to single band (SB) sequences. We also had two secondary hypotheses: first, since ME-ICA denoising removes TE-independent noise, greater activation precision would be observed in multi-echo denoised data (MEdn) relative to ME data without denoising; second, that any MB advantage would be due to the increase in the number of volumes.

Our study focused primarily on univariate effects. However, multivariate analysis techniques, which exploit variance and covariance between voxels and can accommodate participant-specific differences in activation patterns (Coutanche, 2013; Davis et al., 2014; Davis & Poldrack, 2013) are rapidly gaining popularity. These methods frequently rely on the assumption of good-quality signal across the whole brain (Frisby et al., 2023) and so we conducted an exploratory analysis, following the method of Haxby et al. (2001), to decode task condition from the data acquired with each sequence. Finally, we tested for the presence of slice leakage artefacts in our MB data.

To summarise, our study aimed to compare pTx, ME and MB as methods for improving sensitivity in ventral temporal and orbitofrontal regions while maintaining image quality across the rest of the brain.

## 2. Methods

### 2.1. Participants

20 healthy native speakers of British English (age range 18-50, mean age 33.45 years, 12 female, 8 male) participated in the study. All were right-handed, had normal or corrected-to-normal vision, and had no neurological or sensory disorders. All participants gave written informed consent. The research was approved by a local National Health Service (NHS) ethics committee (04/Q105/66).

### 2.2. Stimuli and task

All participants performed a semantic association task and a visual pattern matching task (hereafter called the “control task”) adapted from a previous study (Jung et al., 2017). Each stimulus consisted of three pictures presented simultaneously (Figure 1). Some pictures were line drawings taken from the Pyramids and Palm Trees Test (Howard & Patterson, 1992) and some were colour cartoons or photographs taken from the Camel and Cactus Test (Bozeat et al., 2000). In the semantic task, participants were instructed to indicate which of the two pictures at the bottom of the screen had the closest semantic relationship to the picture at the top (hereafter the “probe picture”). In the control task, participants were instructed to indicate which of two scrambled pictures (generated from the pictures used in the semantic task) at the bottom of the screen was identical to a scrambled probe. There were 248 unique picture triplets, so some stimuli were repeated between runs, but no stimulus was repeated within a run. E-Prime software (Psychology Software Tools Inc., Pittsburgh, USA) was used to display stimuli and record responses. Stimuli were rear-projected onto a screen at the back of the MRI scanner, and observers viewed stimuli through a mirror mounted to the head coil directly above the eyes.

**Figure 1:**
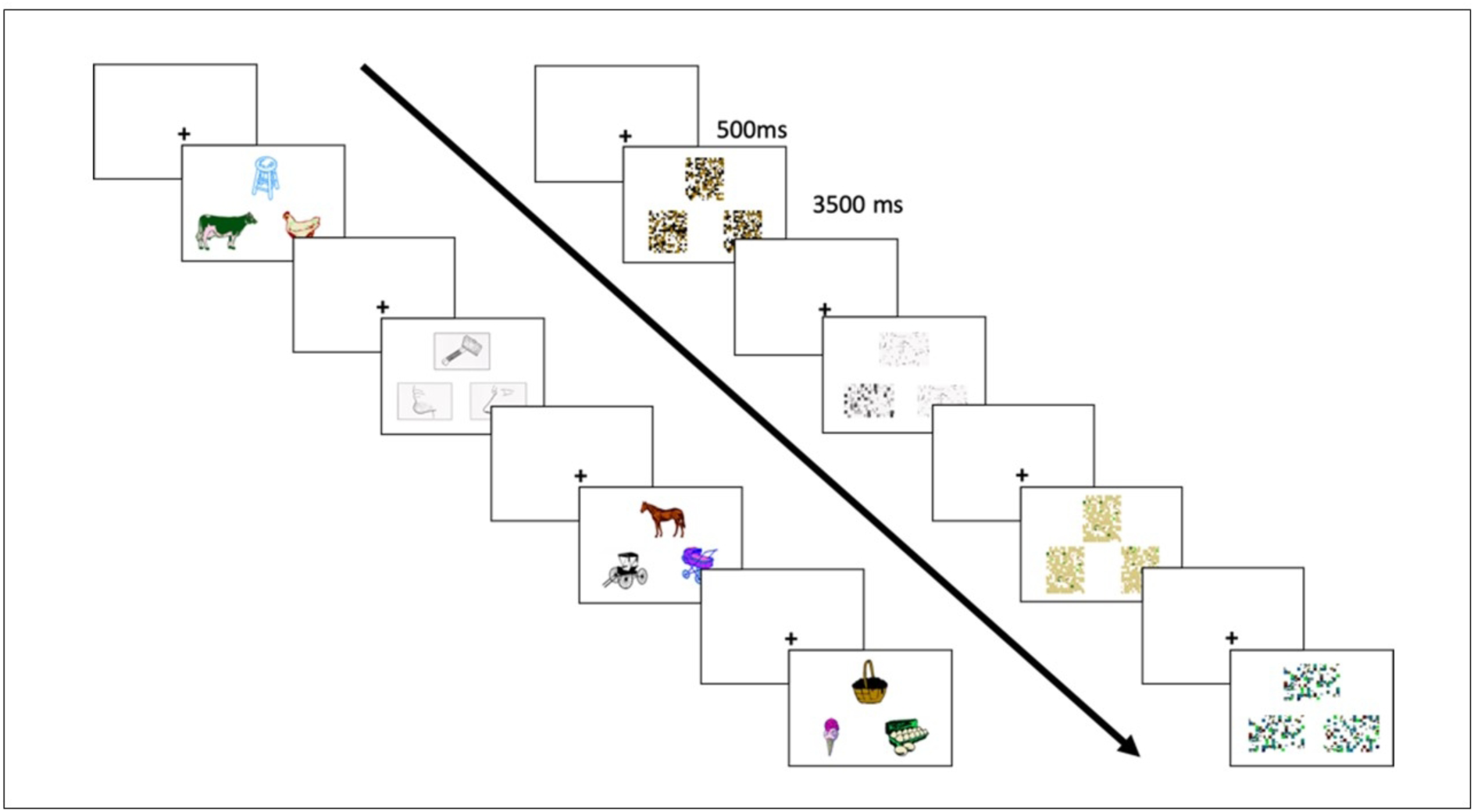
One block of the semantic association task (left of the arrow) and one block of the control task (right of the arrow). Each trial consisted of a fixation cross (500 ms) followed by three pictures presented simultaneously. Participants had to select which of the two pictures at the bottom was more semantically similar to (in the semantic task) or visually identical to (in the control task) the picture at the top (the probe picture). Each block lasted 16 s in total. Reprinted with permission from (Halai et al., 2024).

The study had a block design with three types of block: semantic, control and rest. Each task block consisted of four trials. Each trial consisted of a fixation cross presented for 500 ms followed by a stimulus presented for 3500 ms. Each rest block consisted of a fixation cross presented for 16 s. Each run began 16 s after the start of the MR sequence and then consisted of 30 blocks presented in the order: semantic, control, semantic, control, rest. There were five runs per participant, collected in a single session. Accuracy and reaction time were measured for each trial. Since reaction time is not normally distributed, both metrics were compared across tasks using Wilcoxon’s signed-rank tests and across sequences using Friedman’s nonparametric ANOVA.

### 2.3. Image acquisition

All images were acquired on a whole-body MAGNETOM Terra 7T MRI (Siemens Healthcare, Germany). An 8Tx32Rx head coil (Nova Medical, USA) was used to run all structural and functional imaging sequences.

An MP2RAGE anatomical scan was acquired with the following parameters: 224 sagittal slices (interleaved acquisition), FOV 240 x 225.12 x 240 mm^3^, voxel size 0.75 mm isotropic, TR 4300 ms, inversion times 840 and 2370 ms, TE 1.99 ms, nominal flip angles 5° and 6°, GRAPPA acceleration factor 3, and duration 8 minutes 50 seconds.

Next, manual B_0_-shimming was performed over the volume to be used for EPI. This was necessary because of a known bug in the scanner software which renders automatic shims inaccurate. A double-echo steady state (DESS) sequence was used to generate a field map and the radiographer adjusted shim parameters manually with the aim of reducing the water linewidth, defined as full width of the spectrum at half height, to below 40 Hz. However, for some participants the adjustments proved time-consuming and so adjustment time was capped at 30 minutes from the start of scanning after which the best shim parameters were adopted. The actual average water linewidth was 42 Hz (standard deviation = 9 Hz; missing data for 4 participants). Then a dummy pTx-EPI scan was acquired to trigger the acquisition of subject-specific B_0_ and per-channel B_1_^+^ field maps. Parameters of this dummy scan were the same as the pTx functional sequence (see Table 1 and below) except that only one volume was acquired. The brain was divided into 5 slabs along the slice direction and slab-specific 2-spoke pTx excitation pulses were designed offline. Variable-rate selective excitation (VERSE; Hargreaves et al., 2004) was applied to reduce specific absorption rate (SAR) for the pTx sequence only.

**Table 1:**
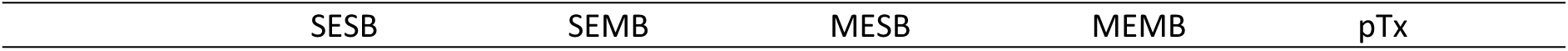
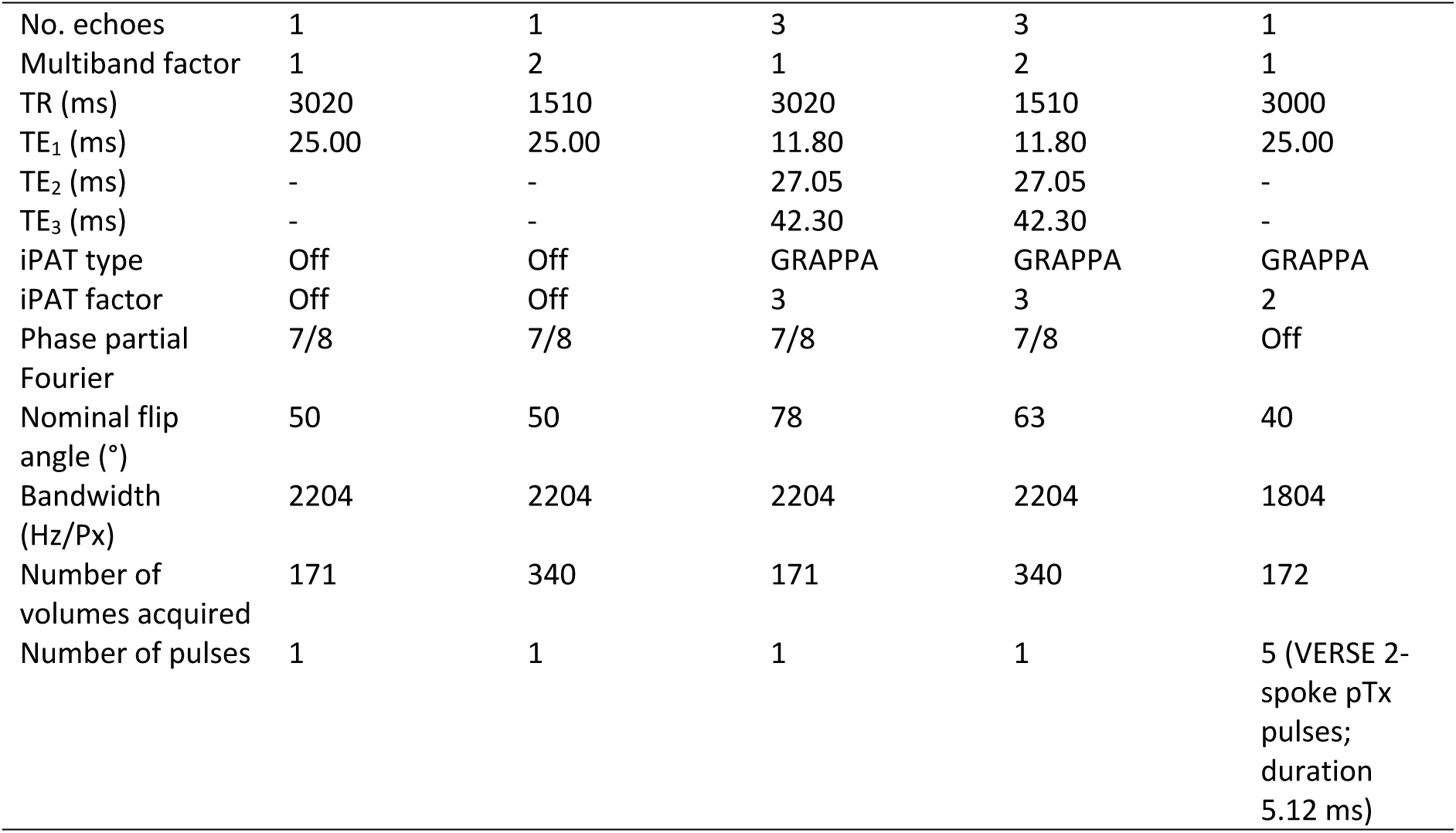
parameters of each sequence.

There were five functional runs of EPI, one run of each sequence. Key parameters are given in Table 1. To ensure that our results would be of use to future cognitive neuroimaging studies, the sequences were designed to represent the way that each parameter would be used in a typical study. Three echoes were used because this is the minimum number needed for ME-ICA and because our scanner hardware prevented us from using four (Halai et al., 2024). An MB factor of 2 was used because previous work suggested that this MB factor, combined with a GRAPPA factor of 3, would be unlikely to cause slice leakage (Todd et al., 2016). GRAPPA and phase partial Fourier were used to ensure a reasonable TE and TR for ME sequences and to ensure that the SE and pTx sequences would have comparable TRs. However, in-plane acceleration reduces tSNR (Yun & Shah, 2017) and using *both* GRAPPA and phase partial Fourier is not needed to achieve a reasonable TE for a single-echo sequence. To facilitate a fair comparison between ME and SE sequences as they would be used in typical studies, no GRAPPA was used for SE sequences. 2 spokes were used because, as the number of spokes increases, the pulse length increases and the bandwidth decreases, and using 2 spokes keeps both those parameters within a reasonable range. 5 slabs were used because, as the number of slabs increases, the time needed for pulse calculation also increases, and using 5 slabs keeps the calculation time short (∼ 5 minutes; Ding et al., 2022). Due to experimenter error, the flip angle was not set to the Ernst angle for SE sequences. The possible impact of this error on results is explored in Supplementary Material 2. Bandwidth of the pTx sequence was selected to match previous work (Ding et al., 2022). It was necessary to increase the bandwidth for our other sequences in order to control TE, but doing so reduces tSNR. To facilitate a fair comparison of pTx to the other sequences as they would be used in typical studies, we did not lower the bandwidth of the pTx sequence to match the others.

The order of sequences was counterbalanced across participants. The following parameters were held constant across sequences: 48 axial slices (interleaved acquisition), FOV 210 x 210 x 210 mm^3^ to cover the whole brain in most participants (visual inspection ensured that the ventral anterior temporal lobe was included in the FOV for all participants and the FOV was tilted up at the nose to avoid ghosting of the eyes into the temporal lobe), voxel size 2.5 mm isotropic (no gap), and A-P phase encoding direction. After each run, 5 further volumes were acquired with the phase encoding direction changed to P-A to facilitate distortion correction during preprocessing.

### 2.4. Data analysis

All analysis code is available at https://github.com/slfrisby/7TOptimisation/.

#### 2.4.1. Preprocessing

For ease of data sharing we converted all DICOMs to BIDS format (Gorgolewski et al., 2016) using *heudiconv* v1.0.0 (Halchenko et al., 2024).

Standard reproducible preprocessing pipelines designed for 3T-fMRI, such as *fMRIprep* (Esteban et al., 2019; Gorgolewski et al., 2011; Markiewicz et al., 2024) perform poorly on 7T-fMRI EPI data. Therefore, the analysis pipeline was split into stages using different software packages.

Since it is notoriously difficult to perform a good-quality brain extraction on MP2RAGE data (because of salt-and-pepper noise in the background and cavities), two pipelines were used. The MP2RAGE T1w (combined) image first had its background noise removed using O’Brien regularisation (O’Brien et al., 2014) and was then submitted to the CAT12 pipeline for segmentation (Gaser et al., 2023; in SPM12; https://www.fil.ion.ucl.ac.uk/spm/). The bias- and global-intensity corrected T1w image produced was provided to as input to the anatomy pipeline in *fMRIPrep* 21.0.1. TheT1w image was skull-stripped with a Nipype implementation of the *antsBrainExtraction.sh* workflow (ANTs 2.3.3; Avants et al., 2009, 2011; https://github.com/ANTsX/ANTs/). Volume-based spatial normalisation to standard space (MNI152NLin2009cAsym) was performed through nonlinear registration with *antsRegistration.sh* (ANTs).

Functional preprocessing was performed using in-house code, composed of functions from AFNI (v.18.3.03; Cox, 1996; Cox & Hyde, 1997; https://afni.nimh.nih.gov/), FSL (v.5.0; Andersson et al., 2003; Jenkinson et al., 2012; Smith, 2002; Smith et al., 2004; https://fsl.fmrib.ox.ac.uk/fsl/fslwiki/), tedana (v.23.0.1; DuPre et al., 2021; Kundu et al., 2011, 2013; The tedana Community et al., 2023; https://tedana.readthedocs.io/en/stable/index.html) and ANTS (v. 2.2.0; Avants et al., 2009, 2011; https://github.com/ANTsX/ANTs/). EPIs were despiked using *3dDespike* (AFNI), slice timing was corrected to the middle slice using *3dTshift* (AFNI), motion was corrected with *3dvolreg* and *3dAllineate* (AFNI) using the first volume of each run as a reference (for ME datasets, the TE_1_ image was aligned and the resulting transform was applied to the TE_2_ and TE_3_ images), and skull-stripped using *BET* (FSL) to create a participant-specific brain mask.

For ME datasets only, *tedana* was used to create two timeseries – one in which the echoes were optimally combined based on T_2_* weighting (Posse, 2012), and one in which the T_2_* optimally-combined data were denoised using ICA. *tedana* conducts denoising by decomposing data using PCA and ICA, classifying components according to whether the signal scales linearly with TE (as the BOLD signal does), and reconstructing the data using only BOLD-like components. The brain mask created with *BET* was used as the mask for this stage.

Next, for all datasets, unwarping was conducted using *topup* and *applytopup* (FSL). Field displacement maps were calculated using ten volumes (five with A-P phase encoding direction, extracted from the start of each functional run, and five with P-A phase encoding direction, collected separately after each run) and the resulting correction was applied to all images. Finally, the mean EPI for each run was coregistered to the skull-stripped native structural image using a rigid-body registration with *AntsRegistrationSyN.sh* (ANTS). EPIs were then transformed into standard space (MNI152NLin2009cAsym) by combining the transforms from native EPI to native T1 and the transforms from native T1 to standard space and applying those transforms to the EPIs using *antsApplyTransforms* (ANTS). Images were smoothed with a 6 mm FWHM Gaussian filter in SPM12 (https://www.fil.ion.ucl.ac.uk/spm/) for GLM analysis.

A separate functional preprocessing pipeline was used to create images for slice leakage analysis. This pipeline differed from the main pipeline in the following ways. Despiking was omitted to avoid the removal of noise. For ME datasets only, rather than run the full *tedana* workflow, we conducted optimal combination of data from multiple echoes (but no denoising) using the *t2smap* command using all voxels in the volume (i.e. no brain mask). All coregistration steps were omitted (the images remained in native EPI space) and no smoothing was applied.

#### 2.4.2. 1^st^-level (within-participant) GLM

Data were analysed using the general linear model (GLM) approach implemented in SPM12 in MATLAB r2019a. We had 5 timeseries of primary interest – standard single-echo single band (SESB), parallel transmit (pTx), single-echo multiband (SEMB), multi-echo single band (MESB) and multi-echo multiband (MEMB). For the latter two sequences, data from all echoes were optimally combined but were not ICA-denoised. We also generated two timeseries with ME-ICA denoising - multi-echo single band with ICA denoising (MESBdn) and multi-echo multiband with ICA denoising (MEMBdn) - and two downsampled MB timeseries created by extracting odd-numbered volumes to match the number of volumes in the single band timeseries - odd-numbered volumes of single-echo multiband (SEMBodd) and odd-volumes of multi-echo multiband (MEMBodd).

At the individual subject level, each block of the semantic and control task was modelled as a boxcar function (resting blocks were modelled implicitly) and these boxcar functions were subsequently convolved with SPM’s difference of gammas haemodynamic response function. The six motion parameters extracted during preprocessing were used as regressors of no interest. The micro-time resolution was set as the number of slices (n = 48), the micro-time onset was set as the reference slice for slice-time correction (n = 24), and the high-pass filter cutoff was 128 seconds. The same MNI template used during preprocessing was used as a mask for the analysis. The parameter estimation method was restricted maximum likelihood estimation (ReML) and serial correlations were accounted for using an autoregressive AR(1) model during estimation. For univariate analyses the contrast of interest was greater activation for the semantic task than the control task (S > C). For exploratory multivariate pattern analysis (MVPA) each block (12 semantic and 12 control) was modelled individually in order to obtain one beta image per block.

Finally, for the slice leakage analysis, the modelling was rerun on the minimally-preprocessed timeseries without any brain mask. We obtained both univariate contrasts of interest (S > C) and beta images per block (for MVPA).

#### 2.4.3. 2^nd^-level (across-participant) GLM

##### 2.4.3.1. Region-of-interest (ROI) analysis

Regions of interest (ROIs) were defined based on a large-scale distortion-corrected 3T-fMRI study of the semantic network (Humphreys et al., 2015). For each comparison, ROIs were analysed only if they overlapped by at least one voxel with the whole-brain contrast of interest (S > C) summed over all sequences in the comparison.

##### 2.4.3.1.1. Univariate analysis

Activation magnitude (1^st^-level beta values) and activation precision (1^st^-level t-values) were extracted from each ROI using a publicly-available script, *roi_extract.m* (https://github.com/MRC-CBU/riksneurotools/blob/master/Util/roi_extract.m). There were two planned t-tests for activation magnitude (pTx > SESB, ME > SE) and four planned t-tests for activation precision (MB > SB, MEdn > ME, MBodd > SB, SB > MBodd). For each planned t-test, results were Bonferroni-corrected for the number of ROIs included.

##### 2.4.3.1.2. Exploratory multivariate pattern analysis (MVPA)

The input to this analysis was beta images, one image per block (12 semantic and 12 control). For each block and each ROI, a vector *v* of beta values was created (voxels within the ROI are numbered arbitrarily):

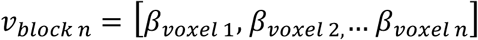

For each ROI, the cosine distance *D* (1 – cosine similarity; (Halai et al., 2024)) between every possible pair of blocks was calculated. For example:

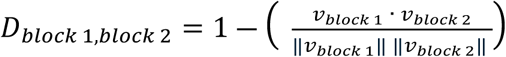

For each ROI, MVPA performance was operationalised as the mean between-task distance (over every possible pair of blocks of different tasks) minus the mean within-task distance (over every possible pair of blocks of the same task). Where block S is a block of the semantic task and block C is a block of the control task, and *n*_*blocks*_ is the total number of blocks of each task (12):

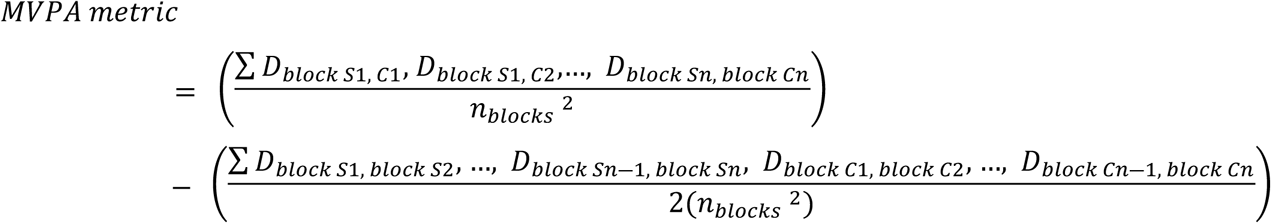

If a region is encoding information about the task, we would expect mean within-task similarity to be large and mean between-task similarity to be small, so the MVPA metric would take a large negative value. If a region is not encoding information about the task, we would expect both within-task and between-task similarity to be zero and the difference between them to be negligible (Halai et al., 2024).

Paired t-tests were used to compare the MVPA metric across sequences (all planned t-tests described in the univariate ROI analysis were conducted; Haxby et al., 2001).

##### 2.4.3.2. Whole-brain analysis

All the above contrasts were assessed at the whole-brain level using t-tests (for comparing pTx and SESB) or using random-effects ANOVAs with one-sample t-tests on the summary statistic (for the three factorial designs: varying echo and band (SESB, SEMB, MESB and MEMB); varying denoising and band (MESBdn, MEMBdn, MESB and MEMB); and varying echo and downsampled band (SEMBodd, MEMBodd, SEMB and MEMB)). The ANOVAs were conducted using a publicly-available script, *batch_spm_anova.m* (https://github.com/MRC-CBU/riksneurotools/blob/master/SPM/batch_spm_anova.m). The group t-maps were assessed for significance by using a voxel-height threshold of p < 0.001 to define clusters and then a cluster-defining family-wise-error corrected threshold of p < 0.05 for statistical inference.

##### 2.4.3.3. Slice leakage analysis

The group-level, whole-brain contrast of interest (S > C) in standard space was inspected and the coordinates of the peak t-values of the top 5 clusters were identified. These coordinates were back-projected to obtain 5 sets of corresponding coordinates in each participant’s native EPI space (hereafter “seeds”, labelled A). Next, voxels to which signal might be warped were identified as artefact locations based on phase shift (FOV/2, labelled B) and, in the MEMB data, GRAPPA (labelled Ag) and phase shift plus GRAPPA (labelled Bg). A spherical ROI, 4 voxels in radius, was defined around each seed location and possible artefact location using a modified version of the scripts developed for McNabb et al. (2020; https://github.com/DrMichaelLindner/MAP4SL/; our version available at https://github.com/slfrisby/7TOptimisation/).

Both univariate (McNabb et al., 2020) and multivariate (Halai et al., 2024) slice leakage tests were conducted. Activation magnitude was extracted from minimally-preprocessed data, and, for the multivariate analysis, the difference between mean within-task similarity and mean between-task similarity was calculated as in the ROI analysis. Paired t-tests were conducted for each seed and artefact location between each sequence of interest (SEMB and MEMB) and a control sequence. For artefact locations based on phase shift (B), the control sequence was the corresponding SB sequence (SESB for SEMB and MESB for MEMB). For artefact locations based on GRAPPA, the control sequence was the corresponding SE sequence, because SE sequences were collected without GRAPPA (SEMB for MEMB). Results were Bonferroni-corrected for the number of peaks (n = 5) and the number of possible artefact regions (n = 1 for SEMB and n = 3 for MEMB).

## 3. Results

### 3.1. Excluded participants

Two participants were excluded because of excessive head motion (this was defined by calculating, for each participant, the percentage of volumes per run with absolute translation values of over 2 mm or absolute rotation values over 1°, averaging these percentages over runs, and excluding any participant whose mean percentage was greater than 2 standard deviations above the mean percentage across participants). One participant was excluded because of technical problems during data acquisition which meant that the pTx run failed. All subsequent analyses were conducted on the remaining 17 participants.

### 3.2. Behavioural results

The 17 participants had good performance on both tasks and, importantly, there were no significant differences between the 5 sequences in terms of accuracy (Friedman’s χ^2^ = 5.26, p = 0.26) or reaction time (Friedman’s χ^2^ = 7.20, p = 0.16). The semantic and control tasks did not differ reliably in accuracy (Wilcoxon’s z = 0; p = 0.06) or reaction time (Wilcoxon’s z = 7; p = 1.00).

### 3.3. Region-of-interest analysis

ROIs that overlapped with the whole-brain contrast of interest (S > C) for at least one of the five sequences are shown in Figure 2.

**Figure 2:**
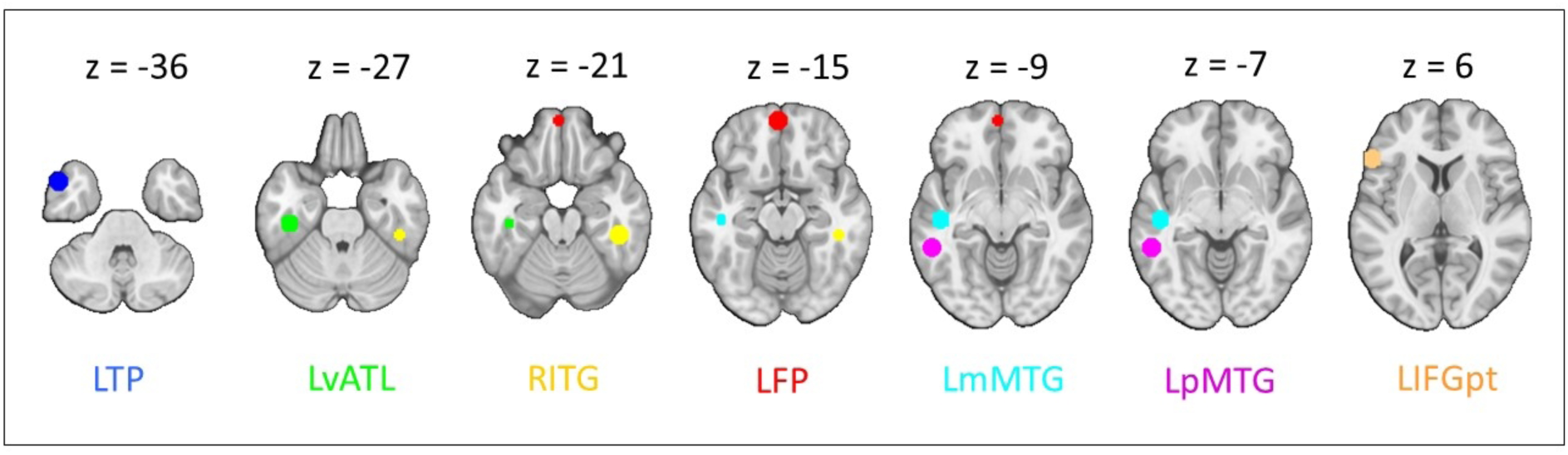
Regions of interest, taken from a meta-analysis of semantic tasks by (Humphreys et al., 2015). All spheres are 8 mm in radius. Regions of interest are overlaid on the MNI152NLin2009cAsym template. LTP = left temporal pole; LvATL = left ventral anterior temporal lobe; RITG = right inferior temporal gyrus; LFP = left frontal pole; LmMTG = left medial middle temporal gyrus; LpMTG = left posterior middle temporal gyrus; LIFGpt = left inferior temporal gyrus pars triangularis.

#### 3.3.1. Univariate analysis

Table 2 shows the p-values for all planned t-contrasts. pTx provided significantly better activation magnitude than SESB in the left posterior middle temporal gyrus (LpMTG). ME sequences provided significantly better activation magnitude than SE sequences in the left ventral anterior temporal lobe (LvATL).

**Table 2:**
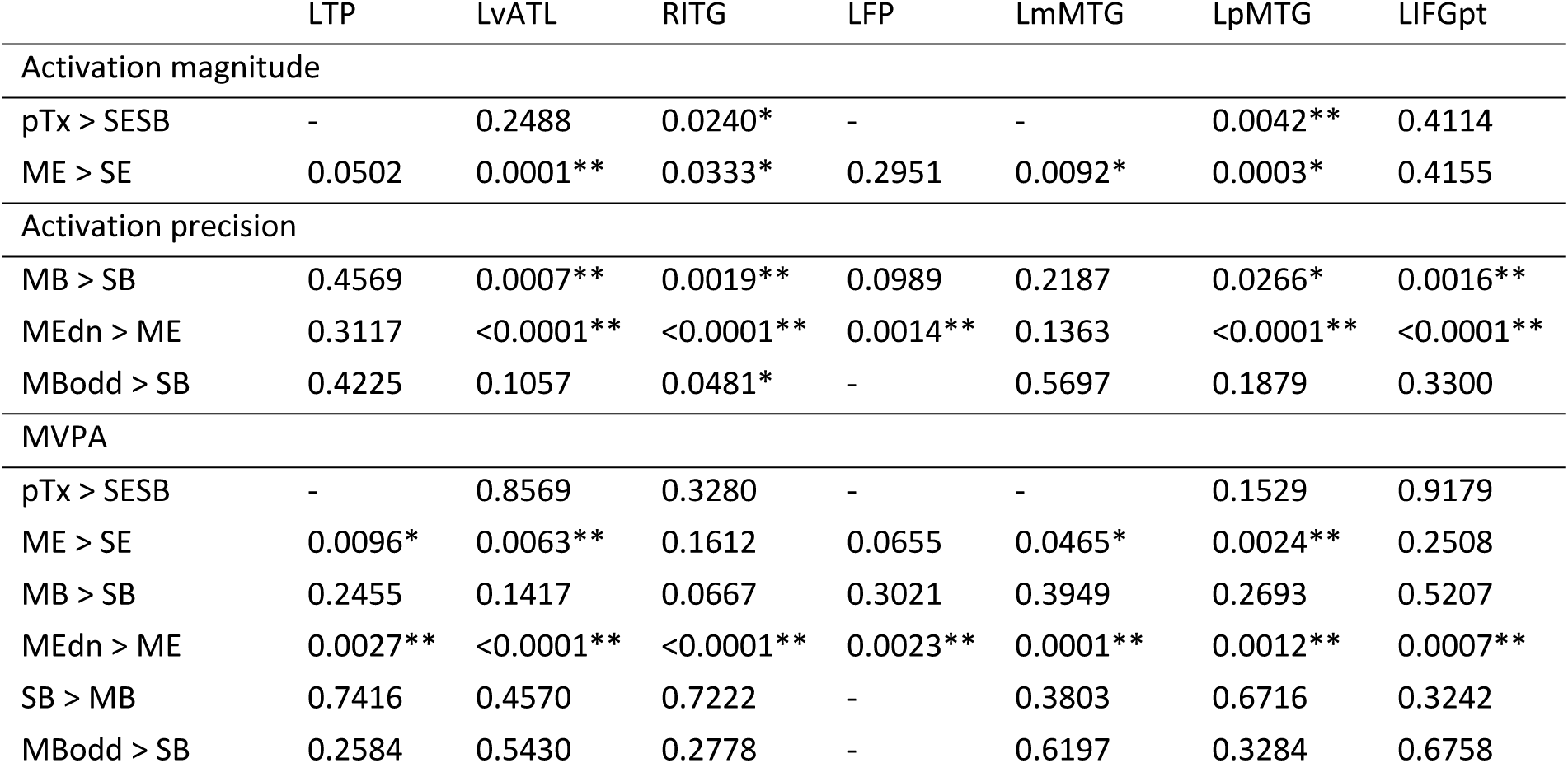
p-values for all t-tests within regions of interest (ROIs) based on the semantic network. * = p < 0.05, ** = p < 0.05 (Bonferroni-corrected for the number of ROIs), - = ROI does not overlap by at least one voxel with the whole-brain contrast of interest (S > C) summed over all sequences in the comparison, LTP = left temporal pole, LvATL = left ventral anterior temporal lobe, RITG = right inferior temporal gyrus, LFP = left frontal pole, LmMTG = left medial middle temporal gyrus, LpMTG = left posterior middle temporal gyrus, LIFGpt = left inferior temporal gyrus pars triangularis.

|  | LTP | LvATL | RITG | LFP | LmMTG | LpMTG | LIFGpt |
| --- | --- | --- | --- | --- | --- | --- | --- |
| Activation magnitude |  |  |  |  |  |  |  |
| pTx > SESB | - | 0.2488 | 0.0240* | - | - | 0.0042** | 0.4114 |
| ME > SE | 0.0502 | 0.0001** | 0.0333* | 0.2951 | 0.0092* | 0.0003* | 0.4155 |
| Activation precision |  |  |  |  |  |  |  |
| MB > SB | 0.4569 | 0.0007** | 0.0019** | 0.0989 | 0.2187 | 0.0266* | 0.0016** |
| MEdn > ME | 0.3117 | <0.0001** | <0.0001** | 0.0014** | 0.1363 | <0.0001** | <0.0001** |
| MBodd > SB | 0.4225 | 0.1057 | 0.0481* | - | 0.5697 | 0.1879 | 0.3300 |
| MVPA |  |  |  |  |  |  |  |
| pTx > SESB | - | 0.8569 | 0.3280 | - | - | 0.1529 | 0.9179 |
| ME > SE | 0.0096* | 0.0063** | 0.1612 | 0.0655 | 0.0465* | 0.0024** | 0.2508 |
| MB > SB | 0.2455 | 0.1417 | 0.0667 | 0.3021 | 0.3949 | 0.2693 | 0.5207 |
| MEdn > ME | 0.0027** | <0.0001** | <0.0001** | 0.0023** | 0.0001** | 0.0012** | 0.0007** |
| SB > MB | 0.7416 | 0.4570 | 0.7222 | - | 0.3803 | 0.6716 | 0.3242 |
| MBodd > SB | 0.2584 | 0.5430 | 0.2778 | - | 0.6197 | 0.3284 | 0.6758 |

MB sequences offered significantly better activation precision than SB sequences in the LvATL, right inferior temporal gyrus (RITG) and left inferior frontal gyrus pars triangularis (LIFGpt). MEdn sequences offered significantly better activation precision than ME in the LvATL, RITG, LpMTG, LIFGpt and left frontal pole (LFP). There was no significant difference in either direction between MBodd sequences and SB sequences (see Supplementary Table 1 for detailed results, plus reverse contrasts). We also found no significant interaction between ME and MB.

#### 3.3.2. Exploratory MVPA

Table 2 also shows the p-values for t-tests comparing our MVPA metric between sequences. The ME sequences produced significantly better performance than SE sequences in the LvATL and LpMTG. MEdn sequences produced better performance than ME sequences in every ROI. All other comparisons failed to reach significance.

### 3.4. Whole-brain analysis

Figure 3 shows the results for activation magnitude. Figure 3A shows a single cluster, within the right fusiform gyrus and lateral inferior occipital cortex, that showed greater activation magnitude with pTx than with SESB (pTx > SESB). Figure 3B shows the main effect of ME over SE, featuring clusters in the fusiform and inferior temporal gyri bilaterally plus the left orbitofrontal cortex (cluster and peak information is provided in Supplementary Table 2).

**Figure 3:**
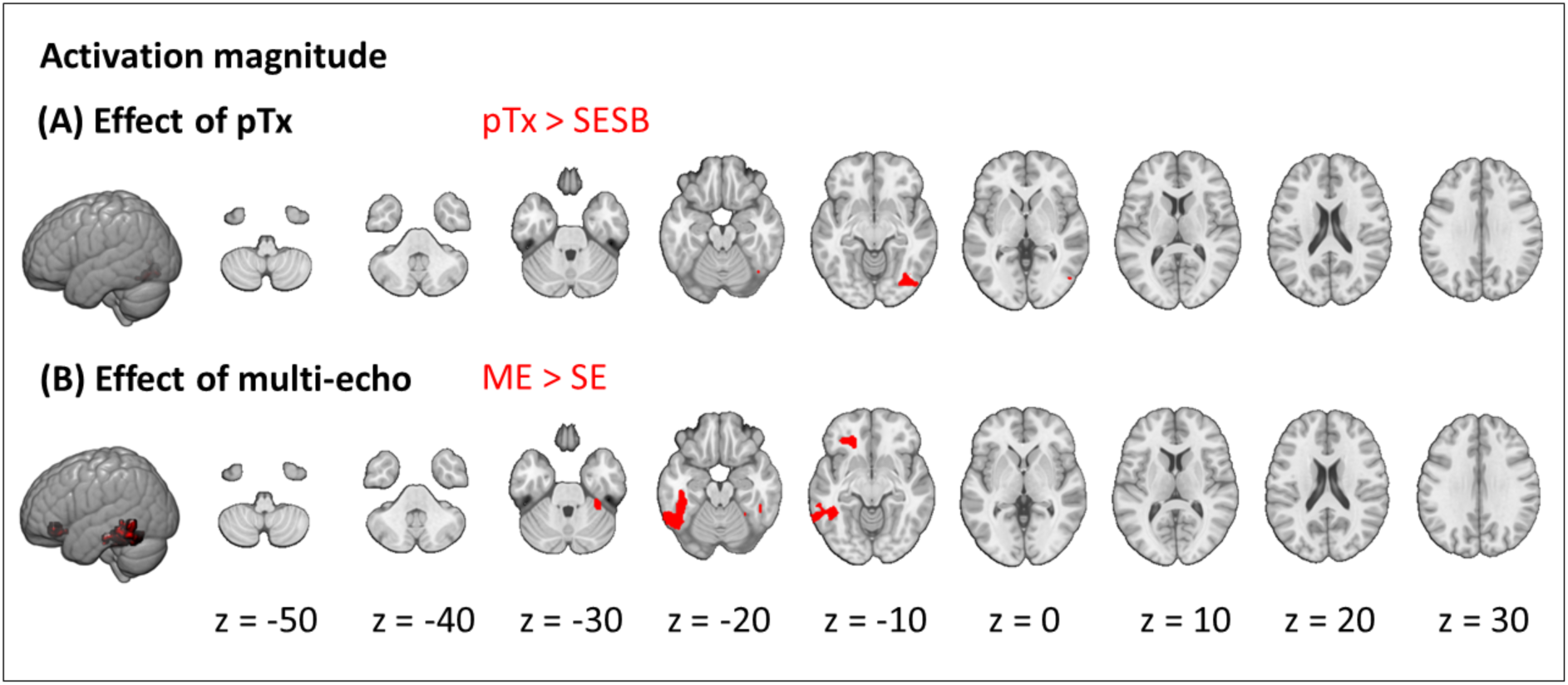
Effects on activation magnitude: (A) effect of parallel transmit (pTx > SESB); (B) effect of echo (ME > SE). Results are cluster-corrected at p < 0.05 based on an uncorrected voxel threshold of p < 0.001 and are overlaid on the MNI152NLin2009cAsym template.

Figure 4 shows the results for activation precision. Both the main effect of MB over SB and the main effect of ME-ICA denoising over ME without ICA denoising extended down the temporal lobe and included frontal regions (Supplementary Table 2). There was no significant difference between downsampled MB sequences and SB sequences in either direction.

**Figure 4:**
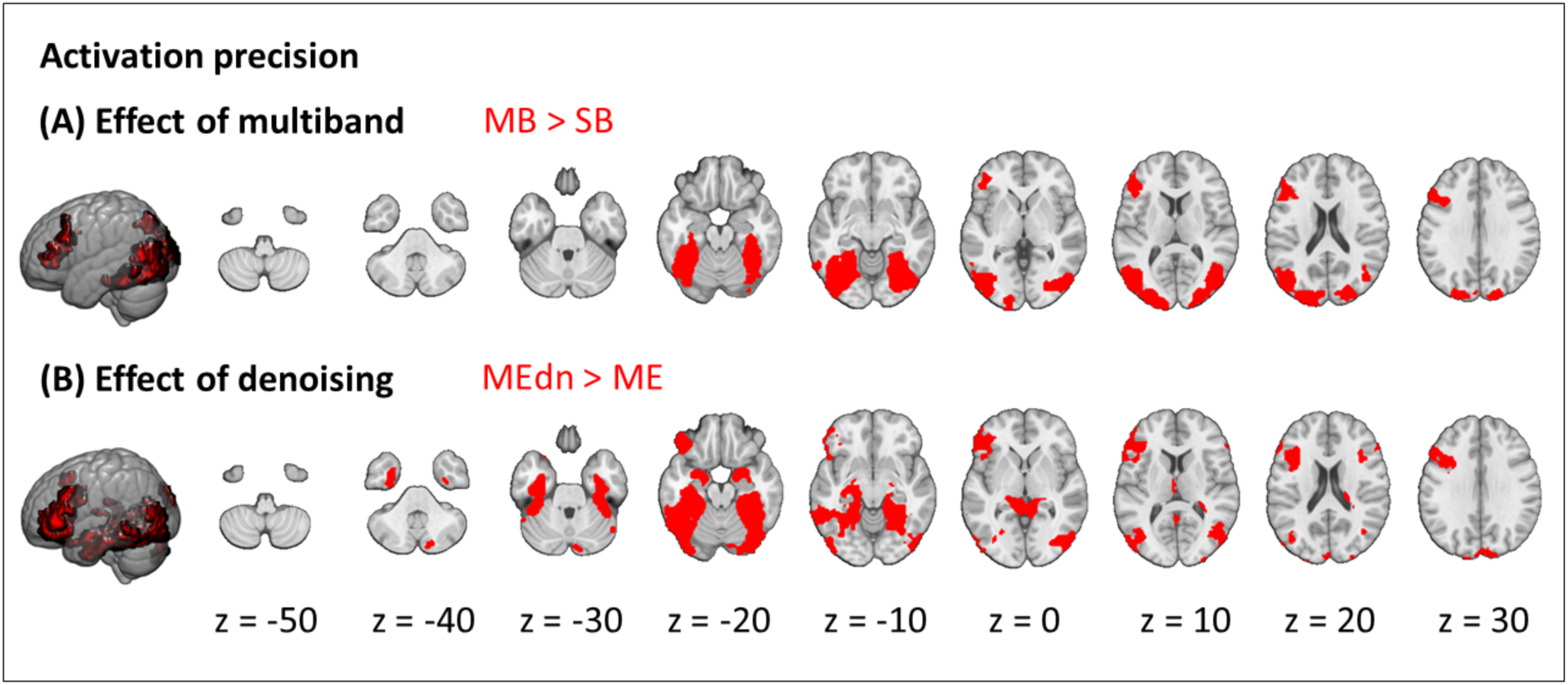
Effects on activation precision: (A) effect of multiband (MB > SB); (B) effect of ME-ICA denoising (MEdn > ME). Results are cluster-corrected at p < 0.05 based on an uncorrected voxel threshold of p < 0.001 and are overlaid on the MNI152NLin2009cAsym template.

To summarise, these results aligned with results from our ROI analyses – ME increased activation magnitude and both MB and ME-ICA denoising increased activation precision (see Supplementary Figures 1-6 for detailed results, plus reverse contrasts).

### 3.5. Slice leakage analysis

Seed and possible artefact locations for an example participant are shown in Figure 5A. Figure 5B shows activation magnitude for possible artefact locations associated with one peak for all participants in each MB sequence and its corresponding sequences. Figure 5C shows MVPA performance. Violin plots for all other peaks are shown in Supplementary Figure 7. t-tests were conducted at each artefact location, but no comparison reached statistical significance (p-values are shown in Supplementary Table 3). t-tests were not conducted at seed locations – if a difference were discovered, this would be attributable to the MB sequence rather than to leakage of signal from the seed into a different location.

**Figure 5:**
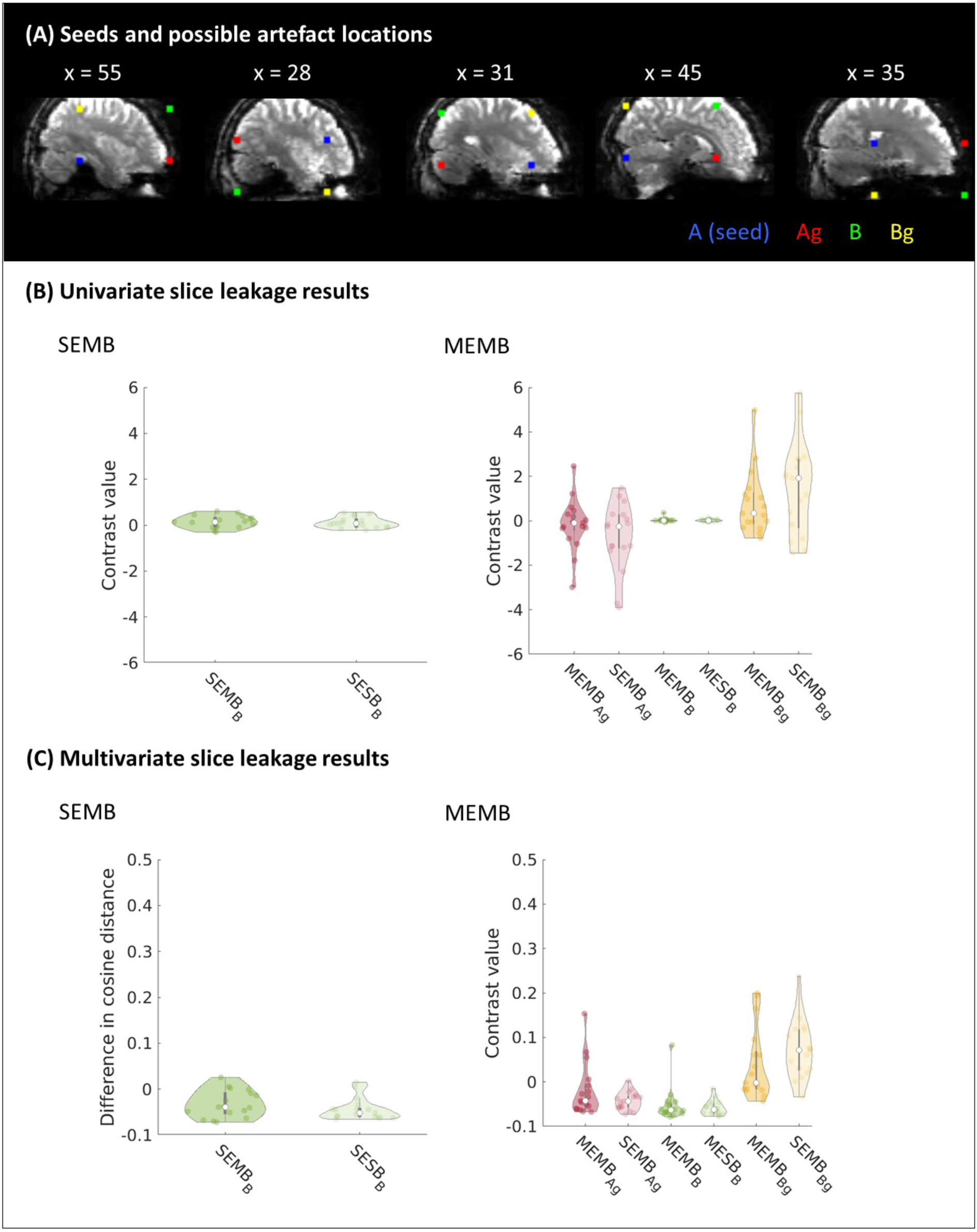
Slice leakage analysis. (A) Seed and possible artefact locations for a single participant. The ROIs (radius 4 voxels) indicate the seed location (blue), possible artefact location based on phase shift (green), possible artefact location based on GRAPPA (red), and possible artefact location based on phase shift and GRAPPA (yellow) in native EPI space. Informed consent was obtained from the participant for this image to be published. (B) Group mean activation magnitude within each possible artefact location corresponding to the first seed for each MB sequence and the corresponding control sequences (plots for other seeds are shown in Supplementary Figure 7; statistics for all seeds are shown in Supplementary Table 3). Each point within a violin plot indicates the mean for one participant (n = 17). (C) Group mean MVPA distance within each sphere for each MB sequence and the corresponding control sequences.

We therefore concluded that there was no evidence for slice leakage in either of our MB datasets.

## 4. Discussion

The tSNR of fMRI varies across the brain. This is especially evident in ventral anterior temporal and orbitofrontal regions, which are located next to the air-filled sinuses and are therefore affected by both B_0_ and B_1_^+^ magnetic field inhomogeneity (Devlin et al., 2000; Gras et al., 2019; Halai et al., 2014, 2015, 2024; Uğurbil, 2018; van der Kolk et al., 2013; Wu et al., 2018). This study is the first 7T-fMRI study systematically comparing pTx, ME and MB as practical methods for improving sensitivity in these regions while maintaining sensitivity across the brain. We found that pTx improved activation magnitude in posterior temporal and occipital regions. ME, however, resulted in improved activation magnitude extending down the temporal lobe and including inferior frontal regions. Both MB and ME-ICA denoising resulted in improved activation precision in the same areas. In an exploratory analysis we found that ME and ME-ICA improved MVPA performance but MB did not. No slice leakage artefacts were associated with our MB sequences.

Although pTx produced better activation magnitude in posterior temporal and occipital regions than the baseline (SESB) sequence without compromising activation magnitude elsewhere, activation magnitude in anterior temporal areas was comparable for both sequences. These results replicated those of Ding et al. (2022) who failed to find improved task contrast in anterior temporal regions with pTx in spite of improved tSNR in the resting state. Note that, despite visible effects of magnetic field inhomogeneity on the EPIs, even the baseline sequence was able to identify semantic activity extending into ventral anterior temporal regions. This is shown in Figure 6 in which the contour of the whole-brain contrast of interest is overlaid on the mean EPI across all participants (detailed contrast maps are shown in Supplementary Figures 1 and 3). It is possible that this finding reflects the increased sensitivity of 7T-fMRI compared to 3T-fMRI, where semantic activity is rarely observed without using a method such as ME (Halai et al., 2014, 2015, 2024) or spin-echo (Binney et al., 2010; Embleton et al., 2010). That a standard sequence can detect signal in these regions might come as a surprise to many neuroimaging researchers as 7T-fMRI is not only associated with increased sensitivity, but also with exacerbated magnetic susceptibility artefacts. This finding should therefore embolden researchers to use 7T fMRI for experimental designs that are difficult to conduct at a lower field strength - for example, studies of special populations, who may benefit from shorter scan times (Cope et al., 2023), or studies requiring high spatial specificity (Marques & Norris, 2018). We do acknowledge that the way we have utilised 7T fMRI in this study will not be suitable for, for example, studies that require ultra-high resolution (in our case, scanner hardware constrained voxel size when combined with a short first echo time).

**Figure 6:**
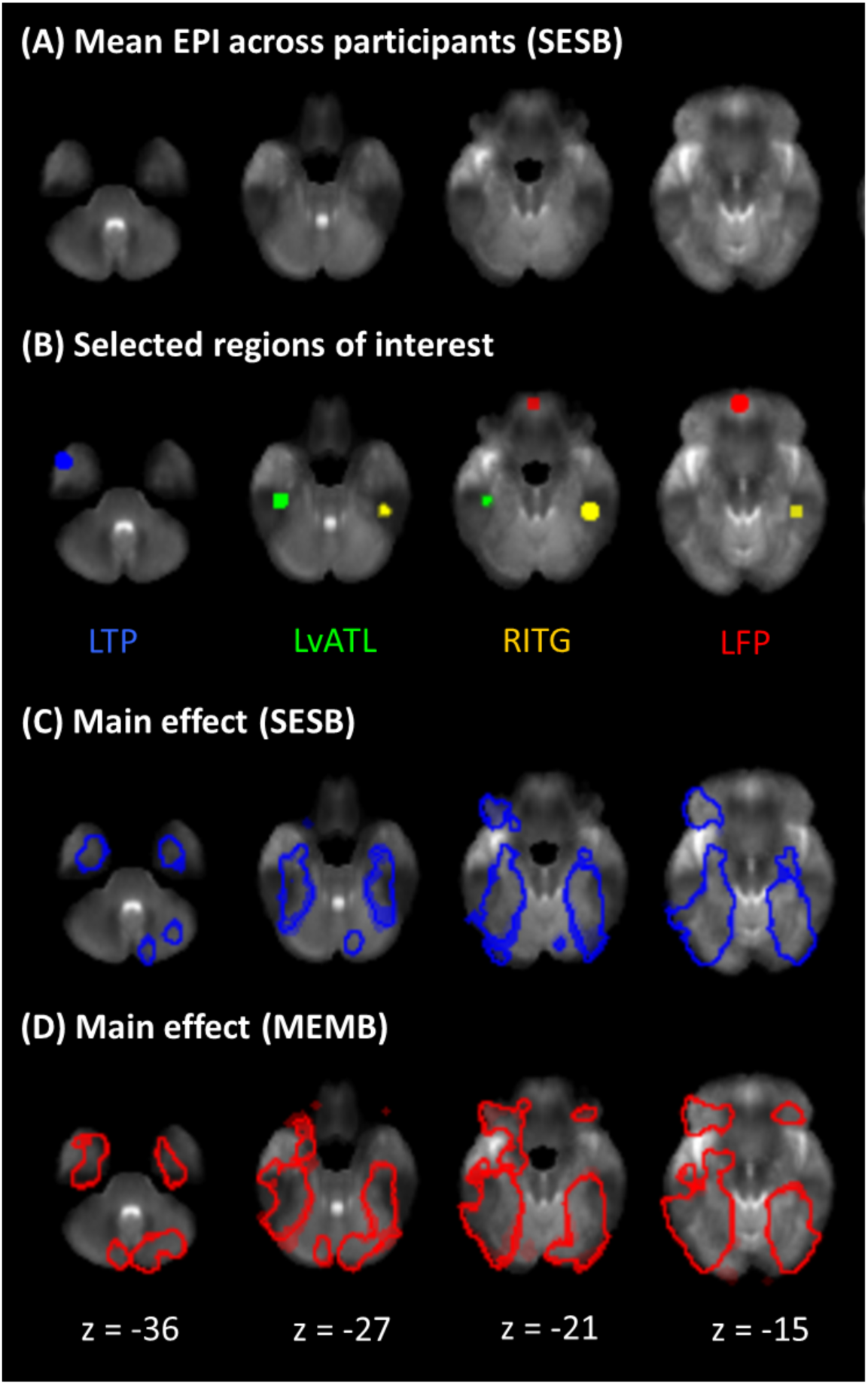
Contrast despite magnetic field inhomogeneity. (A) Mean EPI across all participants for the baseline sequence (SESB) registered to MNI152NLin2009cAsym space. (B) Selected regions of interest overlaid on the mean EPI image. (C) Contour of the contrast of interest (S > C) for the baseline sequence. (D) Contour of the contrast of interest (S > C) for the MEMB sequence. Results are cluster-corrected at p < 0.05 based on an uncorrected voxel threshold of p < 0.001.

The factorial design enabled us to disentangle the effects of ME and MB. As hypothesised, ME improved activation magnitude and these effects were localised to inferior temporal and orbitofrontal areas. Adverse effects on activation magnitude were negligible - although ME reduced activation precision in two very small clusters (Supplementary Figure 4), these were not within areas that the semantic task is known to recruit (Jung et al., 2017). These results demonstrated that having one very short echo is a successful method of increasing activation magnitude in areas where T_2_* is particularly short (Halai et al., 2014, 2015, 2024; Jung et al., 2017). ME also opens up the opportunity for sophisticated denoising such as *tedana* (DuPre et al., 2021; Kundu et al., 2011, 2013; The tedana Community et al., 2023), which improved activation precision in agreement with previous findings (Amemiya et al., 2019; Gonzalez-Castillo et al., 2016). Small clusters in which ME-ICA increased activation magnitude (Supplementary Figure 5) or decreased activation precision (Supplementary Figure 6) were not within regions recruited by the task (Jung et al., 2017).

As hypothesised, MB improved activation precision in areas of inhomogeneity and in agreement with other work (Puckett et al., 2018). Supplementary Figure 4 shows two very small clusters in which MB reduced activation precision, but which lie outside regions recruited by the task (Jung et al., 2017). After downsampling the MB timeseries to match the number of volumes in the SB timeseries, there was no longer any difference between the SB and MB sequences; this suggests that it is the increase in the number of volumes that accounts for the benefits of MB. Additionally, there was no evidence for signal leakage into the simultaneously-acquired slice location (McNabb et al., 2020; Todd et al., 2016), at least for our modest multiband factor of 2. We note that MB and ME seem to have independent effects as the interaction was not significant within any ROI. However, ME seems to offer larger activation magnitude (Figure 6) and MB offers better activation precision. MB can help reduce the longer TR associated with ME and may decrease noise aliasing in studies using short TRs (∼1s).

Although this study focused primarily on univariate effects, we also identified improvements to multivariate metrics that will be of interest to researchers using increasingly-popular MVPA tools (Frisby et al., 2023). ME was found to improve MVPA performance compared to SE sequences and, in turn, MEdn sequences performed significantly better than ME sequences without denoising in every region of interest (with no detrimental effects). Future studies employing multivariate methods should strongly consider using both ME acquisitions and ME-ICA preprocessing.

This study provides evidence supporting use of ME and/or MB sequences to detect signal in regions prone to susceptibility artefacts. It should be noted that, due to experimenter error, the flip angles for the SE sequences were not set to the Ernst angle, which could in theory have accounted for the ME benefit that we observed. However, acquisition of additional SE data with the flip angle set to the Ernst angle indicated that, although use of a suboptimal flip angle compromised tSNR, there was no consistent effect on any of our univariate or multivariate effects of interest (methods and results of this investigation are described in detail in Supplementary Material 2). From this we conclude that the difference in flip angle is not sufficient to explain the ME advantage, although future studies should seek to validate these results using the Ernst angle for all sequences.

It is also important to note that individual 7T MRI scanners have their own hardware and software limitations; this means that each site must optimise and test feasible parameters locally. For example, our system did not allow us to take advantage of the higher spatial resolution offered by 7T-fMRI while retaining an adequately short first echo for our ME sequences. Puckett et al. (2018) used comparable voxel sizes but had a shorter first TE than us (9.9 ms vs our 11.8 ms); Miletić et al. (2020) had higher resolution (1.6 mm vs our 2.5 mm isotropic) and a shorter first echo of 9.66 ms. While we could in principle have achieved a smaller voxel size for our SE sequences, this would have been accompanied either by a narrower FOV or, if we were to maintain the FOV by increasing the number of slices, by a longer TR (which we wished to keep constant across sequences). Additionally, our 7T Terra pTx system imposes very conservative SAR limits (1W per channel, 8W total) for pTx scans but permits 20W total (i.e. 2.5W per channel) for circularly polarised mode (CP-mode or “TrueForm”) scans (equivalent to single transmit). We therefore had to use VERSE-modified pTx pulses which are known to be more sensitive to B_0_ inhomogeneity. We also acknowledge that pTx can be combined with ME or MB (Ding et al., 2023; Wu et al., 2016). Although we used the pTx head coil for all scans, we were not able to create sequences that combined pTx with ME or MB because the software to support this was not implemented on our scanner at the time of running this study. Future studies should seek to test the potential benefits of these combinations. It is possible that studies seeking to test more extreme parameter manipulations (numbers of echoes, multiband factors, or pTx approaches) than we have employed may reach different conclusions to us. For example, Todd et al. (2016) demonstrated that, although multiband factors of 3 or greater can sometimes be associated with greater BOLD sensitivity than the multiband factor of 2 that we have used, higher multiband factors are also associated with a risk of slice leakage. In summary, rather than identifying the limits of what is possible (as others have done), we aim to offer an accessible framework that will enable researchers to leverage 7T-fMRI for investigating the functional roles of regions affected by susceptibility artefacts despite the hardware and software constraints of individual scanners.

## 5. Conclusion

In this study we compared pTx, ME and MB as methods of improving sensitivity in temporal and frontal areas prone to magnetic susceptibility artefacts. Both pTx and ME improved activation magnitude, but only ME showed improvements in artefact-prone regions. MB and ME-ICA improved activation precision in these areas. Exploratory results suggested that both ME and ME-ICA may also benefit MVPA. We demonstrated that a multi-echo, multiband sequence running on the 8Tx32Rx pTx head coil (in CP mode) can detect signal in susceptible regions while maintaining sensitivity across the whole brain and is therefore a versatile choice for future studies using high field strength and investigating the functional roles of ventral temporal and/or orbitofrontal cortex (Binney et al., 2016; Borghesani et al., 2016; Devereux et al., 2018; Duncan, 2010; DuPre et al., 2016; Fernandez et al., 2017; Lambon Ralph et al., 2017; Zahn et al., 2007).

## Acknowledgements

We thank the participants and the Wolfson Brain Imaging Centre radiographers for their assistance.

## Data and code availability

Data will be made publicly available upon peer review and acceptance. Code is publicly available at: https://github.com/slfrisby/7TOptimisation/.

## CRediT statement

Saskia L. Frisby: Conceptualisation, Methodology, Investigation, Formal Analysis, Writing – Original Draft, Writing – Review & Editing, Visualisation; Marta M. Correia: Conceptualisation, Methodology, Writing – Review & Editing, Funding Acquisition; Minghao Zhang: Methodology, Writing – Review & Editing; Christopher T. Rodgers: Methodology, Writing – Review & Editing, Supervision; Timothy T. Rogers: Writing – Review & Editing, Supervision; Matthew A. Lambon Ralph: Writing – Review & Editing, Supervision, Funding Acquisition; Ajay D. Halai: Conceptualisation, Methodology, Formal Analysis, Writing – Review & Editing, Supervision, Funding Acquisition.

## Funding

This work was supported by an MRC Unit grant (SUAG/019 G116768) to M.M.C., an MRC PhD studentship (MR N013433-1) to M.Z., an MRC programme grant (MR/R023883/1) and intramural funding (MC_UU_00005/18) to M.A.L.R, and an MRC Career Development Award (MR/V031481/1) to A.D.H. This work was also supported by the NIHR Cambridge Biomedical Research Centre (NIHR203312) and an MRC Clinical Research Infrastructure Award for 7T (MR/M008983/1). The views expressed are those of the authors and not necessarily those of the NIHR or the Department of Health and Social Care.

## Competing interests

C.T.R. receives research support from Siemens.

## Notes

### Summary of Updates

Incorporating feedback following peer review. Manuscript edited throughout and new supplementary materials added.

