## Supplementary Material 1 for "Optimising 7T-fMRI for imaging regions of magnetic susceptibility"

**Supplementary Material 1: Additional tables and figures**

| | LTP | LvATL | RITG | LFP | LmMTG | LpMTG | LIFGpt |
|---|---|---|---|---|---|---|---|
| Activation magnitude | | | | | | | |
| pTx > SESB | - | 0.2488 | 0.0240* | - | - | 0.0042** | 0.4114 |
| SESB > pTx | - | 0.7512 | 0.9760 | - | - | 0.9958 | 0.5886 |
| ME > SE | 0.0502 | 0.0001** | 0.0333* | 0.2951 | 0.0092* | 0.0003** | 0.4155 |
| SE > ME | 0.9498 | 0.9999 | 0.9667 | 0.7049 | 0.9908 | 0.9997 | 0.5845 |
| MB > SB | 0.4565 | 0.1448 | 0.1124 | 0.3495 | 0.2929 | 0.3023 | 0.3795 |
| SB > MB | 0.5435 | 0.8552 | 0.8876 | 0.6505 | 0.7071 | 0.6977 | 0.6205 |
| Interaction (ME and MB) | 0.8990 | 0.7861 | 0.9897 | 0.7884 | 0.4673 | 0.0569 | 0.9101 |
| MEdn > ME | 0.9609 | 0.9792 | 0.9443 | 0.0783 | 0.5108 | 0.6483 | 0.7065 |
| ME > MEdn | 0.0391* | 0.0208* | 0.0557 | 0.9217 | 0.4892 | 0.3517 | 0.2935 |
| Interaction (MEdn and ME) | 0.2570 | 0.4409 | 0.3560 | 0.3710 | 0.7106 | 0.9530 | 0.3930 |
| MBodd > SB | 0.5085 | 0.2744 | 0.0777 | - | 0.4092 | 0.3466 | 0.4490 |
| SB > MBodd | 0.4915 | 0.7256 | 0.9223 | - | 0.5908 | 0.6534 | 0.5510 |
| Interaction (MBodd and SB) | 0.8817 | 0.8582 | 0.9969 | - | 0.7534 | 0.1315 | 0.8875 |
| Activation precision | | | | | | | |
| pTx > SESB | - | 0.4303 | 0.0191* | - | - | 0.0053** | 0.3239 |
| SESB > pTx | - | 0.5969 | 0.9809 | - | - | 0.9947 | 0.6761 |
| ME > SE | 0.0320* | 0.0017** | 0.0018** | 0.4773 | 0.0027** | <0.0001** | 0.0802 |
| SE > ME | 0.9680 | 0.9983 | 0.9982 | 0.5227 | 0.9973 | 1.0000 | 0.9198 |
| MB > SB | 0.4569 | 0.0007** | 0.0019** | 0.0989 | 0.2187 | 0.0266* | 0.0016** |
| SB > MB | 0.5431 | 0.9993 | 0.9981 | 0.9011 | 0.7813 | 0.9734 | 0.9984 |
| Interaction (ME and MB) | 0.6026 | 0.8131 | 0.9958 | 0.7969 | 0.4791 | 0.0121* | 0.6575 |
| MEdn > ME | 0.3117 | <0.0001** | <0.0001** | 0.0014** | 0.1363 | <0.0001** | <0.0001** |
| ME > MEdn | 0.6883 | 1.0000 | 1.0000 | 0.9986 | 0.8637 | 1.0000 | 1.0000 |
| Interaction (MEdn and ME) | 0.4263 | 0.8147 | 0.9537 | 0.4887 | 0.5175 | 0.6156 | 0.5693 |
| MBodd > SB | 0.4225 | 0.1057 | 0.0481* | - | 0.5697 | 0.1879 | 0.3300 |
| SB > MBodd | 0.5775 | 0.8943 | 0.9519 | - | 0.4303 | 0.8121 | 0.6700 |
| Interaction (MBodd and SB) | 0.7872 | 0.9387 | 1.0000 | - | 0.8476 | 0.0915 | 0.7912 |
| MVPA | | | | | | | |
| pTx > SESB | - | 0.8569 | 0.3280 | - | - | 0.1529 | 0.9179 |
| SESB > pTx | - | 0.1431 | 0.6720 | - | - | 0.8471 | 0.0821 |
| ME > SE | 0.0096* | 0.0063** | 0.1612 | 0.0655 | 0.0465* | 0.0024** | 0.2508 |
| SE > ME | 0.9904 | 0.9937 | 0.8388 | 0.9345 | 0.9535 | 0.9976 | 0.7492 |
| MB > SB | 0.2455 | 0.1417 | 0.0667 | 0.3021 | 0.3949 | 0.2693 | 0.5207 |
| SB > MB | 0.7076 | 0.8460 | 0.9351 | 0.6933 | 0.5945 | 0.6776 | 0.4744 |
| Interaction (ME and MB) | 0.6612 | 0.6182 | 0.9791 | 0.1553 | 0.1998 | 0.1660 | 0.8216 |
| MEdn > ME | 0.0027** | <0.0001** | <0.0001** | 0.0023** | <0.0001** | 0.0012** | 0.0007** |

| | | | | | | | |
|---|---|---|---|---|---|---|---|
| ME > MEdn | 0.9973 | 1.0000 | 1.0000 | 0.9977 | 0.9999 | 0.9988 | 0.9993 |
| Interaction (MEdn and ME) | 0.2711 | 0.8973 | 0.8733 | 0.3296 | 0.8462 | 0.9714 | 0.9560 |
| MBodd > SB | 0.2584 | 0.5430 | 0.2778 | - | 0.6197 | 0.3284 | 0.6758 |
| SB > MBodd | 0.7416 | 0.4570 | 0.7222 | - | 0.3803 | 0.6716 | 0.3242 |
| Interaction (MBodd and SB) | 0.8113 | 0.8003 | 0.9960 | - | 0.2954 | 0.1939 | 0.8072 |

Supplementary table 1: p-values for all statistical tests within regions of interest (ROIs) based on the semantic network. * = p < 0.05, ** = p < 0.05 (Bonferroni-corrected for the number of ROIs), - = ROI does not overlap by at least one voxel with the whole-brain contrast of interest (S > C) summed over all sequences in the comparison, LTP = left temporal pole, LvATL = left ventral anterior temporal lobe, RITG = right inferior temporal gyrus, LFP = left frontal pole, LmMTG = left medial middle temporal gyrus, LpMTG = left posterior middle temporal gyrus, LIFGpt = left inferior temporal gyrus pars triangularis.

| | Cluster extent (voxels) | z-value | x | y | z | Anatomical label |
|---|---|---|---|---|---|---|
| **Activation magnitude** | | | | | | |
| Comparing pTx and SESB | | | | | | |
| SESB | 8941 | 7.55 | -29 | -42 | -13 | Left temporal fusiform cortex pos |
| | | 7.39 | 34 | -47 | -20 | Right temporal occipital fusiform cortex |
| | | 7.16 | -31 | -34 | -23 | Left temporal fusiform cortex pos |
| | | 6.93 | -39 | -52 | -18 | Left temporal occipital fusiform cortex |
| | | 6.83 | 31 | -57 | -18 | Right temporal occipital fusiform cortex |
| | | 6.76 | 26 | -42 | -18 | Right temporal occipital fusiform cortex |
| | | 6.71 | -49 | -77 | 17 | Left lateral occipital cortex sup |
| | | 6.57 | -41 | -74 | 4 | Left lateral occipital cortex inf |
| | 2294 | 6.75 | -46 | 43 | -3 | Left frontal pole |
| | | 6.73 | -54 | 40 | -8 | Left frontal pole |
| | | 6.72 | -49 | 33 | 10 | Left inferior frontal gyrus p tri |
| | | 6.43 | -44 | 20 | 20 | Left inferior frontal gyrus p ope |
| | | 6.18 | -51 | 36 | 24 | Left frontal pole |
| | | 6.13 | -39 | 6 | 34 | Left middle frontal gyrus |
| | | 5.94 | -36 | 16 | 24 | Left inferior frontal gyrus p ope |

| | | | | | | |
|---|---|---|---|---|---|---|
| | | 5.5 | -41 | 8 | 22 | Left inferior frontal gyrus p ope |
| | 232 | 5.91 | -6 | 33 | 44 | Left superior frontal gyrus |
| | 182 | 5.59 | -26 | 16 | 42 | Left middle frontal gyrus |
| | | 3.6 | -39 | 20 | 57 | Left middle frontal gyrus |
| | 160 | 5.01 | 41 | 23 | 20 | Right middle frontal gyrus |
| | | 4.27 | 36 | 16 | 30 | Right middle frontal gyrus |
| | 66 | 4.3 | 41 | 33 | -8 | Right frontal orbital cortex |
| | 110 | 4.21 | -6 | -60 | 17 | Left precuneous cortex |
| | | 4.1 | -4 | -52 | 12 | Left precuneous cortex |
| | | 3.89 | -11 | -57 | 7 | Left precuneous cortex |
| pTx | 11682 | 7.79 | -29 | -42 | -13 | Left temporal fusiform cortex pos |
| | | 7.69 | 34 | -47 | -20 | Right temporal occipital fusiform cortex |
| | | 7.52 | 36 | -67 | -16 | Right occipital fusiform gyrus |
| | | 7.52 | -39 | -52 | -18 | Left temporal occipital fusiform cortex |
| | | 7.43 | -36 | -42 | -20 | Left temporal fusiform cortex pos |
| | | 7.34 | 41 | -42 | -18 | Right temporal occipital fusiform cortex |
| | | 7.3 | -29 | -32 | -20 | Left parahippocampal gyrus pos |
| | | 7.17 | 46 | -77 | -8 | Right lateral occipital cortex inf |
| | 3242 | 6.79 | -54 | 40 | -8 | Left frontal pole |
| | | 6.72 | -51 | 36 | 24 | Left frontal pole |
| | | 6.67 | -46 | 43 | -3 | Left frontal pole |
| | | 6.67 | -49 | 33 | 10 | Left inferior frontal gyrus p tri |
| | | 6.63 | -39 | 6 | 34 | Left middle frontal gyrus |
| | | 6.59 | -44 | 20 | 20 | Left inferior frontal gyrus p ope |
| | | 6.22 | -51 | 20 | 30 | Left middle frontal gyrus |
| | | 5.83 | -29 | 18 | 62 | Left superior frontal gyrus |
| | 501 | 6.1 | 14 | -74 | -30 | Right cerebellum |
| | | 6.06 | 9 | -82 | -30 | Right cerebellum |
| | | 5.34 | 34 | -74 | -48 | Right cerebellum |
| | 296 | 5.57 | 44 | 26 | 20 | Right inferior frontal gyrus p tri |
| | | 4.59 | 36 | 16 | 30 | Right middle frontal gyrus |

| | | | | | | |
|---|---|---|---|---|---|---|
| | | 3.28 | 49 | 33 | 34 | Right middle frontal gyrus |
| | 97 | 4.61 | 24 | -60 | 20 | Right precuneous cortex |
| | | 4.03 | 19 | -50 | 10 | Right precuneous cortex |
| | | 4.01 | 11 | -52 | 14 | Right precuneous cortex |
| | 146 | 4.4 | 34 | 38 | -13 | Right frontal pole |
| | | 4.2 | 31 | 33 | -20 | Right frontal pole |
| | | 3.63 | 34 | 30 | 0 | Right frontal orbital cortex |
| | | 3.53 | 51 | 33 | -16 | Right frontal orbital cortex |
| | | 3.43 | 46 | 26 | -8 | Right frontal orbital cortex |
| ANOVA | 126 | 4.62 | 36 | -67 | -16 | Right occipital fusiform gyrus |
| | | 4.08 | 39 | -77 | -10 | Right lateral occipital cortex inf |
| | | 3.9 | 31 | -80 | -13 | Right occipital fusiform gyrus |
| | | 3.39 | 51 | -72 | 0 | Right lateral occipital cortex inf |
| pTx > SESB | 172 | 4.76 | 36 | -67 | -16 | Right occipital fusiform gyrus |
| | | 4.24 | 39 | -77 | -10 | Right lateral occipital cortex inf |
| | | 4.07 | 31 | -80 | -13 | Right occipital fusiform gyrus |
| | | 3.57 | 51 | -72 | 0 | Right lateral occipital cortex inf |
| | | 3.18 | 54 | -70 | 7 | Right lateral occipital cortex inf |
| 2 x 2 factorial design – echo and band | | | | | | |
| SESB | 11459 | Inf | -31 | -44 | -20 | Left temporal fusiform cortex pos |
| | | Inf | 34 | -47 | -20 | Right temporal occipital fusiform cortex |
| | | Inf | 36 | -40 | -23 | Right temporal occipital fusiform cortex |
| | | Inf | -39 | -57 | -16 | Left temporal occipital fusiform cortex |
| | | Inf | 46 | -80 | 0 | Right lateral occipital cortex inf |
| | | Inf | -44 | -77 | 4 | Left lateral occipital cortex inf |
| | | Inf | 49 | -67 | 12 | Right lateral occipital cortex inf |
| | | Inf | 31 | -44 | -13 | Right temporal occipital fusiform cortex |
| | 2394 | Inf | -49 | 33 | 10 | Left inferior frontal gyrus p tri |
| | | 7.68 | -39 | 10 | 24 | Left inferior frontal gyrus p ope |
| | | 7.25 | -46 | 20 | 22 | Left inferior frontal gyrus p ope |
| | | 7.2 | -46 | 43 | -3 | Left frontal pole |

| | | | | | | |
|---|---|---|---|---|---|---|
| | | 7.12 | -49 | 33 | -3 | Left inferior frontal gyrus p tri |
| | | 6.84 | -44 | 40 | -10 | Left frontal pole |
| | | 6.66 | -56 | 18 | 27 | Left inferior frontal gyrus p ope |
| | | 6.62 | -59 | 23 | 14 | Left inferior frontal gyrus p tri |
| | 304 | 7.68 | 39 | 20 | 20 | Right inferior frontal gyrus p ope |
| | | 4.78 | 54 | 30 | 12 | Right inferior frontal gyrus p tri |
| | 276 | 6.44 | -6 | 23 | 50 | Left superior frontal gyrus |
| | | 5.12 | -6 | 33 | 44 | Left superior frontal gyrus |
| | 206 | 5.89 | -29 | 20 | 52 | Left middle frontal gyrus |
| | | 4.09 | -39 | 20 | 57 | Left middle frontal gyrus |
| | | 3.78 | -34 | 10 | 64 | Left middle frontal gyrus |
| | 140 | 5.4 | 34 | -70 | -46 | Right cerebellum |
| | | 3.49 | 24 | -77 | -43 | Right cerebellum |
| | 109 | 4.95 | 11 | -77 | -28 | Right cerebellum |
| | | 4.06 | 14 | -80 | -36 | Right cerebellum |
| | 113 | 4.4 | -9 | -52 | 14 | Left precuneous cortex |
| | | 4.33 | -11 | -54 | 4 | Left precuneous cortex |
| SEMB | 20421 | Inf | -31 | -44 | -20 | Left temporal fusiform cortex pos |
| | | Inf | 34 | -47 | -20 | Right temporal occipital fusiform cortex |
| | | Inf | 49 | -67 | 12 | Right lateral occipital cortex inf |
| | | Inf | 44 | -77 | -3 | Right lateral occipital cortex inf |
| | | Inf | -39 | -57 | -16 | Left temporal occipital fusiform cortex |
| | | Inf | 31 | -34 | -20 | Right temporal fusiform cortex pos |
| | | Inf | 39 | -40 | -23 | Right temporal occipital fusiform cortex |
| | | Inf | 36 | -64 | -16 | Right occipital fusiform gyrus |
| | 568 | 7.7 | 39 | 20 | 17 | Right inferior frontal gyrus p ope |
| | | 7.46 | 49 | 28 | 17 | Right inferior frontal gyrus p tri |
| | | 6.37 | 39 | 16 | 30 | Right middle frontal gyrus |
| | | 3.95 | 19 | 10 | 32 | Right lateral ventricle |
| | 225 | 5.46 | 44 | 30 | -6 | Right frontal orbital cortex |
| | | 4.57 | 34 | 33 | -18 | Right frontal pole |

| | | | | | | |
|---|---|---|---|---|---|---|
| | 203 | 5.09 | -14 | 3 | 10 | left anterior thalamic radiation |
| | | 5.05 | -9 | -12 | 4 | left thalamus |
| MESB | 22950 | Inf | -31 | -44 | -20 | Left temporal fusiform cortex pos |
| | | Inf | 34 | -47 | -20 | Right temporal occipital fusiform cortex |
| | | Inf | -44 | -50 | -16 | Left inferior temporal gyrus temocc |
| | | Inf | -39 | -57 | -18 | Left temporal occipital fusiform cortex |
| | | Inf | 36 | -67 | -16 | Right occipital fusiform gyrus |
| | | Inf | 34 | -60 | -18 | Right temporal occipital fusiform cortex |
| | | Inf | -41 | -40 | -18 | Left temporal fusiform cortex pos |
| | | Inf | -34 | -70 | -16 | Left occipital fusiform gyrus |
| | 733 | Inf | 41 | 20 | 22 | Right inferior frontal gyrus p ope |
| | | Inf | 49 | 28 | 20 | Right inferior frontal gyrus p tri |
| | | 6.23 | 54 | 33 | 14 | Right inferior frontal gyrus p tri |
| | | 4.57 | 54 | 30 | 4 | Right inferior frontal gyrus p tri |
| | 468 | 7.75 | 31 | 36 | -13 | Right frontal pole |
| | | 5.91 | 39 | 28 | -3 | Right frontal orbital cortex |
| | 212 | 6.95 | -11 | -82 | -33 | Left cerebellum |
| MEMB | 20648 | Inf | -34 | -44 | -23 | Left temporal fusiform cortex pos |
| | | Inf | 36 | -42 | -23 | Right temporal occipital fusiform cortex |
| | | Inf | -36 | -57 | -16 | Left temporal occipital fusiform cortex |
| | | Inf | -44 | -50 | -16 | Left inferior temporal gyrus temocc |
| | | Inf | 34 | -60 | -18 | Right temporal occipital fusiform cortex |
| | | Inf | 36 | -67 | -16 | Right occipital fusiform gyrus |
| | | Inf | -34 | -70 | -16 | Left occipital fusiform gyrus |
| | | Inf | -44 | -40 | -18 | Left temporal fusiform cortex pos |
| | 461 | 7.65 | 41 | 23 | 20 | Right middle frontal gyrus |
| | | 7.24 | 49 | 28 | 20 | Right inferior frontal gyrus p tri |
| | 234 | 6.64 | 31 | 36 | -13 | Right frontal pole |
| | | 4.67 | 41 | 30 | -6 | Right frontal orbital cortex |
| | 167 | 6.09 | -9 | -84 | -33 | Left cerebellum |

| | | | | | | |
|---|---|---|---|---|---|---|
| ANOVA | 268 | 5.3 | -44 | -40 | -18 | Left temporal fusiform cortex pos |
| | | 4.71 | -51 | -57 | -20 | Left inferior temporal gyrus temocc |
| | | 4.39 | -59 | -50 | -16 | Left inferior temporal gyrus temocc |
| | | 4.09 | -36 | -44 | -26 | Left temporal occipital fusiform cortex |
| | | 3.77 | -41 | -47 | -8 | left inferior longitudinal fas |
| | | 3.63 | -66 | -54 | -6 | Left middle temporal gyrus temocc |
| | 108 | 4.44 | -34 | 18 | 20 | Left inferior frontal gyrus p ope |
| | | 4.09 | -36 | 18 | 32 | Left middle frontal gyrus |
| | | 3.61 | -44 | 26 | 22 | Left inferior frontal gyrus p tri |
| | | 3.33 | -39 | 26 | 14 | Left inferior frontal gyrus p tri |
| | 80 | 4.31 | 41 | -54 | 17 | Right angular gyrus |
| | | 4.08 | 46 | -67 | 10 | Right lateral occipital cortex inf |
| | | 3.64 | 29 | -60 | 0 | Right lingual gyrus |
| | | 3.62 | 34 | -70 | 4 | right inferior longitudinal fas |
| | 123 | 4.27 | 26 | -47 | -3 | Right lingual gyrus |
| | | 4.18 | 34 | -44 | -16 | Right temporal occipital fusiform cortex |
| | | 3.97 | 24 | -32 | -13 | right cingulum hipp |
| | | 3.86 | 31 | -24 | -20 | Right parahippocampal gyrus pos |
| | | 3.5 | 34 | -34 | -20 | Right temporal fusiform cortex pos |
| ME > SE | 620 | 5.98 | -44 | -40 | -18 | Left temporal fusiform cortex pos |
| | | 5.27 | -51 | -57 | -20 | Left inferior temporal gyrus temocc |
| | | 5.12 | -59 | -50 | -16 | Left inferior temporal gyrus temocc |
| | | 4.6 | -36 | -44 | -26 | Left temporal occipital fusiform cortex |
| | | 4.51 | -41 | -47 | -10 | left inferior longitudinal fas |
| | | 4.43 | -66 | -54 | -6 | Left middle temporal gyrus temocc |
| | | 4.18 | -44 | -24 | -20 | Left inferior temporal gyrus post |
| | | 3.53 | -36 | -24 | -26 | Left temporal fusiform cortex pos |
| | 101 | 5.44 | 34 | -40 | -28 | Right temporal fusiform cortex pos |
| | | 3.55 | 46 | -40 | -20 | Right inferior temporal gyrus temocc |
| | 117 | 5.11 | -21 | 36 | -10 | Left frontal orbital cortex |
| | | 3.68 | -34 | 36 | -10 | Left frontal orbital cortex |
| | | 3.43 | -9 | 23 | -13 | Left subcallosal cortex |

| | | | | | | |
|---|---|---|---|---|---|---|
| 2 x 2 factorial design – denoising and band | | | | | | |
| MEdn > ME | 323 | 4.62 | -31 | -47 | 2 | Left lateral ventricle |
| | | 4.22 | -14 | -34 | 4 | left thalamus |
| | | 4.13 | -29 | -37 | 14 | left anterior thalamic radiation |
| | | 4.08 | 4 | -32 | 12 | 3rd ventricle |
| | | 4.08 | -16 | -42 | 17 | Left lateral ventricle |
| | | 4.05 | 11 | -32 | 4 | right thalamus |
| | 104 | 4.55 | -1 | -52 | -56 | Left cerebellum |
| | | 3.75 | -11 | -40 | -50 | brain stem |
| | | 3.59 | 4 | -54 | -66 | brain stem |
| | 135 | 3.94 | 29 | -34 | 4 | Right lateral ventricle |
| | | 3.81 | 29 | -34 | 17 | Right lateral ventricle |
| | | 3.79 | 21 | -24 | 22 | right corticospinal tract |
| | | 3.64 | 31 | -44 | 0 | Right lateral ventricle |
| 2 x 2 factorial design – echo and (downsampled) band | | | | | | |
| ANOVA | 224 | 4.77 | -49 | -52 | -20 | Left inferior temporal gyrus temocc |
| | | 4.72 | -44 | -40 | -18 | Left temporal fusiform cortex pos |
| | | 4.02 | -41 | -47 | -8 | left inferior longitudinal fas |
| | | 4.02 | -59 | -50 | -16 | Left inferior temporal gyrus temocc |
| | | 3.68 | -39 | -44 | -26 | Left temporal fusiform cortex pos |
| | 85 | 4.34 | 41 | -57 | 17 | Right angular gyrus |
| | | 4.18 | 44 | -67 | 10 | Right lateral occipital cortex inf |
| | 89 | 4.19 | -34 | 13 | 20 | Left inferior frontal gyrus p ope |
| | | 4.04 | -36 | 18 | 32 | Left middle frontal gyrus |
| | | 3.56 | -44 | 28 | 22 | Left middle frontal gyrus |
| **Activation precision** | | | | | | |
| Comparing pTx and SESB | | | | | | |
| SESB | 9906 | 7.62 | -29 | -42 | -13 | Left temporal fusiform cortex pos |
| | | 7.35 | 39 | -44 | -18 | Right temporal occipital fusiform cortex |
| | | 7.29 | 34 | -52 | -20 | Right temporal occipital fusiform cortex |
| | | 7.18 | -34 | -50 | -20 | Left temporal occipital fusiform cortex |

| | | | | | | |
|---|---|---|---|---|---|---|
| | | 7.18 | 34 | -67 | -16 | Right occipital fusiform gyrus |
| | | 7.06 | 24 | -42 | -18 | Right temporal occipital fusiform cortex |
| | | 7.01 | 36 | -60 | -18 | Right temporal occipital fusiform cortex |
| | | 7 | 51 | -70 | 7 | Right lateral occipital cortex inf |
| | 2162 | 6.27 | -54 | 40 | -8 | Left frontal pole |
| | | 5.97 | -51 | 36 | 10 | Left inferior frontal gyrus p tri |
| | | 5.91 | -41 | 13 | 22 | Left inferior frontal gyrus p ope |
| | | 5.89 | -54 | 30 | 17 | Left inferior frontal gyrus p tri |
| | | 5.68 | -36 | 36 | -13 | Left frontal orbital cortex |
| | | 5.56 | -51 | 20 | 22 | Left inferior frontal gyrus p ope |
| | | 5.36 | -46 | 36 | -16 | Left frontal pole |
| | | 5.3 | -39 | 6 | 34 | Left middle frontal gyrus |
| | 383 | 5.47 | -24 | 16 | 44 | Left superior frontal gyrus |
| | | 5.14 | -9 | 23 | 47 | Left superior frontal gyrus |
| | | 4.97 | -29 | 23 | 57 | Left superior frontal gyrus |
| | | 4.68 | -6 | 30 | 44 | Left superior frontal gyrus |
| | | 3.81 | -14 | 18 | 42 | Left paracingulate gyrus |
| | 165 | 5.43 | 31 | 0 | -36 | Right temporal fusiform cortex ant |
| | | 3.69 | 31 | -10 | -36 | Right temporal fusiform cortex pos |
| | 185 | 5.19 | 44 | 26 | 17 | Right inferior frontal gyrus p tri |
| | 129 | 4.83 | 14 | -74 | -30 | Right cerebellum |
| | | 3.73 | 14 | -90 | -30 | Right cerebellum |
| | 121 | 4.59 | -6 | -60 | 17 | Left precuneous cortex |
| | | 3.7 | -16 | -57 | 10 | Left precuneous cortex |
| | 91 | 4.44 | 36 | -64 | -38 | Right cerebellum |
| | | 3.9 | 34 | -67 | -48 | Right cerebellum |
| | | 3.8 | 29 | -72 | -40 | Right cerebellum |
| pTx | 12474 | 7.73 | 34 | -67 | -16 | Right occipital fusiform gyrus |
| | | 7.7 | 34 | -52 | -20 | Right temporal occipital fusiform cortex |
| | | 7.7 | -29 | -42 | -13 | Left temporal fusiform cortex pos |

| | | | | | |
|---|---|---|---|---|---|
| | 7.68 | 39 | -44 | -18 | Right temporal occipital fusiform cortex |
| | 7.53 | 24 | -42 | -18 | Right temporal occipital fusiform cortex |
| | 7.32 | 36 | -60 | -18 | Right temporal occipital fusiform cortex |
| | 7.26 | -36 | -50 | -20 | Left temporal occipital fusiform cortex |
| | 7.21 | 34 | -37 | -20 | Right temporal fusiform cortex pos |
| 2923 | 6.34 | -54 | 40 | -8 | Left frontal pole |
| | 6.22 | -54 | 30 | 17 | Left inferior frontal gyrus p tri |
| | 6 | -41 | 16 | 22 | Left inferior frontal gyrus p ope |
| | 5.97 | -39 | 6 | 34 | Left middle frontal gyrus |
| | 5.95 | -51 | 40 | 7 | Left frontal pole |
| | 5.92 | -46 | 43 | -3 | Left frontal pole |
| | 5.86 | -46 | 36 | -16 | Left frontal pole |
| | 5.81 | -51 | 38 | 24 | Left frontal pole |
| 333 | 5.82 | 44 | 26 | 17 | Right inferior frontal gyrus p tri |
| | 4.17 | 36 | 16 | 30 | Right middle frontal gyrus |
| 637 | 5.74 | 9 | -84 | -33 | Right cerebellum |
| | 5.44 | 14 | -77 | -30 | Right cerebellum |
| | 5.26 | 34 | -74 | -53 | Right cerebellum |
| | 4.69 | 36 | -70 | -38 | Right cerebellum |
| | 4.26 | 24 | -82 | -48 | Right cerebellum |
| 336 | 5.21 | -6 | 30 | 44 | Left superior frontal gyrus |
| | 4.4 | -6 | 43 | 30 | Left paracingulate gyrus |
| | 4.36 | -1 | 28 | 52 | Left superior frontal gyrus |
| | 3.47 | -11 | 53 | 37 | Left frontal pole |
| 179 | 4.7 | 24 | -60 | 17 | Right supracalcarine cortex |
| | 4.41 | 21 | -57 | 10 | Right precuneous cortex |
| | 4.27 | 9 | -52 | 10 | Right precuneous cortex |
| 159 | 4.43 | 34 | 38 | -16 | Right frontal pole |
| | 4.11 | 51 | 33 | -16 | Right frontal orbital cortex |
| | 4.06 | 34 | 30 | 0 | Right frontal orbital cortex |
| | 3.4 | 44 | 38 | -23 | Right frontal pole |

| 2 x 2 factorial design – echo and band | | | | | | |
|---|---|---|---|---|---|---|
| SESB | 9047 | Inf | -29 | -44 | -18 | Left temporal occipital fusiform cortex |
| | | Inf | 36 | -42 | -20 | Right temporal occipital fusiform cortex |
| | | Inf | -29 | -52 | -10 | Left temporal occipital fusiform cortex |
| | | Inf | 49 | -67 | 12 | Right lateral occipital cortex inf |
| | | Inf | 31 | -60 | -16 | Right temporal occipital fusiform cortex |
| | | 7.7 | 26 | -50 | -13 | Right temporal occipital fusiform cortex |
| | | 7.69 | -46 | -74 | 12 | Left lateral occipital cortex inf |
| | | 7.68 | 34 | -34 | -23 | Right temporal fusiform cortex pos |
| | 1738 | 6.46 | -49 | 33 | 7 | Left inferior frontal gyrus p tri |
| | | 6.37 | -39 | 13 | 22 | Left inferior frontal gyrus p ope |
| | | 6.19 | -44 | 26 | 12 | Left inferior frontal gyrus p tri |
| | | 6.18 | -46 | 20 | 20 | Left inferior frontal gyrus p ope |
| | | 5.13 | -41 | 40 | -10 | Left frontal pole |
| | | 5.12 | -44 | 33 | -13 | Left frontal orbital cortex |
| | | 5.11 | -46 | 46 | 0 | Left frontal pole |
| | | 4.85 | -49 | 38 | -18 | Left frontal pole |
| | 213 | 6.28 | 36 | 20 | 20 | Right inferior frontal gyrus p ope |
| | | 5.53 | 49 | 28 | 17 | Right inferior frontal gyrus p tri |
| | 136 | 5.16 | -6 | 23 | 47 | Left superior frontal gyrus |
| | | 4.04 | -9 | 33 | 44 | Left superior frontal gyrus |
| | 123 | 4.62 | -29 | 20 | 54 | Left middle frontal gyrus |
| SEMB | 22882 | Inf | -29 | -47 | -18 | Left temporal occipital fusiform cortex |
| | | Inf | 46 | -67 | 12 | Right lateral occipital cortex inf |
| | | Inf | 34 | -34 | -23 | Right temporal fusiform cortex pos |
| | | Inf | -46 | -77 | 10 | Left lateral occipital cortex inf |
| | | Inf | 29 | -54 | -13 | Right temporal occipital fusiform cortex |
| | | Inf | 36 | -42 | -20 | Right temporal occipital fusiform cortex |
| | | Inf | -26 | -52 | -10 | Left temporal occipital fusiform cortex |

| | | | | | | |
|---|---|---|---|---|---|---|
| | | Inf | -41 | -80 | -10 | Left lateral occipital cortex inf |
| | 452 | Inf | 51 | 28 | 17 | Right inferior frontal gyrus p tri |
| | | 4.7 | 39 | 16 | 30 | Right middle frontal gyrus |
| | 248 | 5.42 | 34 | 33 | -20 | Right frontal pole |
| | | 5.22 | 44 | 33 | -8 | Right frontal pole |
| | 104 | 5.36 | 6 | -62 | -50 | Right cerebellum |
| | | 4.9 | 9 | -54 | -43 | Right cerebellum |
| | 198 | 5.35 | -6 | 20 | -28 | Left subcallosal cortex |
| | | 3.55 | -4 | 43 | -30 | Left frontal medial cortex |
| | | 3.37 | -11 | 10 | -26 | Left frontal orbital cortex |
| | | 3.32 | 6 | 23 | -26 | Right subcallosal cortex |
| | 210 | 5.35 | -9 | -12 | 7 | left thalamus |
| | | 5.03 | -14 | 3 | 4 | left anterior thalamic radiation |
| | 141 | 5.16 | -9 | -82 | -33 | Right cerebellum |
| | 104 | 5.1 | -9 | 63 | -18 | Left frontal pole |
| MESB | 23065 | Inf | -31 | -44 | -18 | Left temporal occipital fusiform cortex |
| | | Inf | 36 | -42 | -20 | Right temporal occipital fusiform cortex |
| | | Inf | -39 | -42 | -16 | Left temporal fusiform cortex pos |
| | | Inf | -46 | -74 | 10 | Left lateral occipital cortex inf |
| | | Inf | 34 | -34 | -23 | Right temporal fusiform cortex pos |
| | | Inf | -26 | -52 | -13 | Left temporal occipital fusiform cortex |
| | | Inf | 34 | -54 | -18 | Right temporal occipital fusiform cortex |
| | | Inf | -44 | -50 | -13 | Left inferior temporal gyrus temocc |
| | 1169 | 7.74 | 36 | 20 | 20 | Right inferior frontal gyrus p ope |
| | | 7.72 | 49 | 28 | 17 | Right inferior frontal gyrus p tri |
| | | 6.92 | 34 | 36 | -13 | Right frontal pole |
| | | 6.03 | 31 | 10 | 30 | Right middle frontal gyrus |
| | | 5.87 | 41 | 28 | -3 | Right frontal orbital cortex |
| | 216 | 5.84 | -9 | -82 | -33 | Left cerebellum |
| | 130 | 5.24 | 6 | -54 | -43 | Right cerebellum |
| | | 5.01 | 4 | -62 | -50 | Right cerebellum |

| | | | | | | |
|---|---|---|---|---|---|---|
| | | 3.45 | -6 | -60 | -48 | Left cerebellum |
| MEMB | 23862 | Inf | -31 | -44 | -18 | Left temporal occipital fusiform cortex |
| | | Inf | -36 | -52 | -18 | Left temporal occipital fusiform cortex |
| | | Inf | -44 | -77 | 2 | Left lateral occipital cortex inf |
| | | Inf | -49 | -74 | 10 | Left lateral occipital cortex inf |
| | | Inf | 34 | -44 | -20 | Right temporal occipital fusiform cortex |
| | | Inf | -41 | -42 | -18 | Left temporal fusiform cortex pos |
| | | Inf | -31 | -64 | -13 | Left occipital fusiform gyrus |
| | | Inf | -44 | -50 | -13 | Left inferior temporal gyrus temocc |
| | 1016 | Inf | 49 | 28 | 17 | Right inferior frontal gyrus p tri |
| | | 7.76 | 39 | 23 | 17 | Right inferior frontal gyrus p tri |
| | | 6.83 | 34 | 36 | -13 | Right frontal pole |
| | | 6.42 | 36 | 13 | 32 | Right middle frontal gyrus |
| | | 5.55 | 41 | 28 | -3 | Right frontal orbital cortex |
| | | 4.66 | 49 | 33 | 7 | Right frontal pole |
| | 223 | 6.3 | -9 | -84 | -36 | Left cerebellum |
| | 143 | 5.46 | 4 | -57 | -48 | Right cerebellum |
| | | 5.3 | 6 | -54 | -40 | Right cerebellum |
| | | 3.66 | -6 | -57 | -48 | Left cerebellum |
| | 90 | 5.28 | -21 | 68 | 17 | Left frontal pole |
| ANOVA | 2729 | 6.34 | -34 | -44 | -20 | Left temporal fusiform cortex pos |
| | | 6.15 | -44 | -40 | -18 | Left temporal fusiform cortex pos |
| | | 6.14 | -44 | -77 | 2 | Left lateral occipital cortex inf |
| | | 6.07 | -49 | -77 | 10 | Left lateral occipital cortex inf |
| | | 5.99 | -44 | -50 | -16 | Left inferior temporal gyrus temocc |
| | | 5.88 | -36 | -72 | -16 | Left occipital fusiform gyrus |
| | | 5.69 | -41 | -80 | -10 | Left lateral occipital cortex inf |
| | | 5.35 | -51 | -44 | -13 | left superior longitudinal fas |
| | 2159 | 5.73 | 46 | -67 | 12 | Right lateral occipital cortex inf |
| | | 5.64 | 36 | -40 | -28 | Right temporal fusiform cortex pos |
| | | 5.6 | 36 | -74 | -13 | Right occipital fusiform gyrus |
| | | 5.44 | 46 | -77 | -3 | Right lateral occipital cortex inf |

| | | | | | | |
|---|---|---|---|---|---|---|
| | | 5.19 | 29 | -60 | -13 | Right temporal occipital fusiform cortex |
| | | 5.17 | 44 | -32 | -23 | Right inferior temporal gyrus post |
| | | 5.05 | 46 | -77 | 7 | Right lateral occipital cortex inf |
| | | 4.96 | 36 | -52 | -20 | Right temporal occipital fusiform cortex |
| | 188 | 5.01 | -21 | 36 | -10 | Left frontal orbital cortex |
| | | 4.06 | 6 | 6 | -16 | Right subcallosal cortex |
| | | 3.84 | 4 | 13 | -13 | Right subcallosal cortex |
| | | 3.81 | -9 | 26 | -13 | Left subcallosal cortex |
| | | 3.5 | -34 | 33 | -13 | Left frontal orbital cortex |
| | 90 | 5 | 34 | 3 | -38 | Right temporal pole |
| | | 4.13 | 34 | -12 | -30 | Right parahippocampal gyrus ant |
| | | 4.12 | 34 | 8 | -48 | Right temporal pole |
| | 501 | 4.89 | -34 | 13 | 20 | Left inferior frontal gyrus p ope |
| | | 4.68 | -36 | 18 | 30 | Left middle frontal gyrus |
| | | 4.56 | -46 | 28 | 22 | Left inferior frontal gyrus p tri |
| | | 4.31 | -46 | 33 | 7 | Left inferior frontal gyrus p tri |
| | | 4.17 | -54 | 23 | 30 | Left middle frontal gyrus |
| | | 4.06 | -44 | 26 | 10 | Left inferior frontal gyrus p tri |
| | | 3.29 | -56 | 33 | 14 | Left inferior frontal gyrus p tri |
| | 97 | 4.87 | -31 | -74 | 47 | Left lateral occipital cortex sup |
| | 613 | 4.79 | -24 | -94 | 10 | Left occipital pole |
| | | 4.56 | -21 | -94 | 20 | Left occipital pole |
| | | 3.83 | -9 | -90 | 17 | Left occipital pole |
| | | 3.81 | -21 | -90 | 32 | Left occipital pole |
| | 109 | 4.74 | 31 | 36 | -10 | Right frontal pole |
| | | 4.07 | 14 | 30 | -10 | right uncinate fas |
| | 80 | 4.36 | 14 | -82 | -46 | Right cerebellum |
| | | 3.93 | 31 | -70 | -43 | Right cerebellum |
| | | 3.48 | 36 | -77 | -46 | Right cerebellum |
| ME > SE | 1309 | 6.42 | -44 | -40 | -18 | Left temporal fusiform cortex pos |
| | | 5.93 | -51 | -44 | -13 | left superior longitudinal fas |

| | | | | | | |
|---|---|---|---|---|---|---|
| | | 5.69 | -41 | -47 | -8 | left inferior longitudinal fas |
| | | 5.14 | -54 | -57 | -20 | Left inferior temporal gyrus temocc |
| | | 5.01 | -46 | -52 | -20 | Left inferior temporal gyrus temocc |
| | | 4.9 | -56 | -77 | 14 | Left lateral occipital cortex inf |
| | | 4.71 | -46 | -42 | 10 | Left superior temporal gyurs pos |
| | | 4.58 | -39 | -24 | -20 | Left temporal fusiform cortex pos |
| | 348 | 6.13 | 36 | -40 | -28 | Right temporal fusiform cortex pos |
| | | 4.35 | 24 | -52 | -18 | Right temporal occipital fusiform cortex |
| | | 4.19 | 34 | -22 | -36 | Right temporal fusiform cortex pos |
| | | 3.98 | 44 | -32 | -23 | Right inferior temporal gyrus post |
| | | 3.9 | 34 | -12 | -33 | Right temporal fusiform cortex pos |
| | | 3.73 | 46 | -44 | -16 | Right inferior temporal gyrus temocc |
| | | 3.27 | 24 | -44 | -23 | Right temporal occipital fusiform cortex |
| | 562 | 5.51 | -19 | 36 | -13 | Left frontal orbital cortex |
| | | 5.12 | 31 | 38 | -10 | Right frontal pole |
| | | 4.84 | 9 | 8 | -16 | Right subcallosal cortex |
| | | 4.66 | 14 | 30 | -10 | right uncinate fas |
| | | 4.44 | -9 | 23 | -13 | Left subcallosal cortex |
| | | 3.89 | -34 | 36 | -10 | Left frontal orbital cortex |
| | | 3.77 | 1 | 18 | -18 | Right subcallosal cortex |
| | | 3.76 | 11 | 20 | -13 | Right subcallosal cortex |
| | 134 | 4.95 | 14 | -82 | -46 | Right cerebellum |
| | | 3.64 | 26 | -77 | -46 | Right cerebellum |
| | | 3.53 | 11 | -82 | -36 | Right cerebellum |
| | | 3.4 | 14 | -72 | -36 | Right cerebellum |
| | 92 | 4.34 | 54 | -74 | -10 | Right lateral occipital cortex inf |
| | | 4.32 | 49 | -84 | 2 | Right lateral occipital cortex inf |
| | | 4.11 | 54 | -77 | 10 | Right lateral occipital cortex inf |
| SE > ME | 93 | 4.89 | -36 | 10 | 17 | Left frontal operculum cortex |
| | | 3.71 | -34 | -2 | 32 | Left precentral gyrus |
| MB > SB | 3959 | 6.7 | -44 | -77 | 2 | Left lateral occipital cortex inf |

| | | | | | | |
|---|---|---|---|---|---|---|
| | | 6.58 | -49 | -77 | 10 | Left lateral occipital cortex inf |
| | | 6.53 | -34 | -47 | -20 | Left temporal occipital fusiform cortex |
| | | 6.43 | -36 | -72 | -16 | Left occipital fusiform gyrus |
| | | 6.28 | -41 | -80 | -10 | Left lateral occipital cortex inf |
| | | 6.26 | -36 | -54 | -18 | Left temporal occipital fusiform cortex |
| | | 5.95 | -44 | -50 | -16 | Left inferior temporal gyrus temocc |
| | | 5.7 | -29 | -54 | -13 | Left temporal occipital fusiform cortex |
| | 2569 | 6.18 | 46 | -67 | 12 | Right lateral occipital cortex inf |
| | | 6.1 | 46 | -77 | -3 | Right lateral occipital cortex inf |
| | | 6.01 | 36 | -74 | -13 | Right occipital fusiform gyrus |
| | | 5.65 | 31 | -62 | -16 | Right occipital fusiform gyrus |
| | | 5.62 | 46 | -77 | 7 | Right lateral occipital cortex inf |
| | | 5.49 | 29 | -90 | 14 | Right occipital pole |
| | | 5.35 | 36 | -52 | -20 | Right temporal occipital fusiform cortex |
| | | 5.3 | 34 | -34 | -23 | Right temporal fusiform cortex pos |
| | 828 | 4.68 | -54 | 23 | 30 | Left middle frontal gyrus |
| | | 4.62 | -46 | 33 | 12 | Left inferior frontal gyrus p tri |
| | | 4.47 | -46 | 26 | 22 | Left inferior frontal gyrus p tri |
| | | 4.39 | -39 | 16 | 30 | Left middle frontal gyrus |
| | | 4.12 | -51 | 20 | 42 | Left middle frontal gyrus |
| | | 3.59 | -44 | 38 | -6 | Left frontal pole |
| | | 3.51 | -46 | 48 | 7 | Left frontal pole |
| | | 3.44 | -39 | 23 | 52 | Left middle frontal gyrus |
| SB > MB | 427 | 4.36 | 44 | -44 | 57 | Right superior parietal lobule |
| | | 4.2 | 46 | -40 | 42 | Right supramarginal gyrus pos |
| | | 3.65 | 56 | -32 | 50 | Right supramarginal gyrus ant |
| | | 3.64 | 31 | -47 | 64 | Right superior parietal lobule |
| | | 3.53 | 59 | -40 | 52 | Right supramarginal gyrus pos |
| | | 3.49 | 26 | -44 | 57 | Right superior parietal lobule |
| | | 3.35 | 61 | -47 | 37 | Right supramarginal gyrus pos |
| | 130 | 3.95 | -4 | -60 | 64 | Left precuneous cortex |

| | | | | | | |
|---|---|---|---|---|---|---|
| | | 3.59 | -11 | -57 | 74 | Left superior parietal lobule |
| | | 3.56 | 1 | -52 | 67 | Right precuneous cortex |
| | | 3.47 | -21 | -57 | 62 | Left superior parietal lobule |
| | | 3.45 | -4 | -54 | 54 | Left precuneous cortex |
| | | 3.37 | -11 | -64 | 62 | Left lateral occipital cortex sup |
| 2 x 2 factorial design – denoising and band | | | | | | |
| ANOVA | 6202 | Inf | 36 | -32 | -26 | Right temporal fusiform cortex pos |
| | | 6.88 | 39 | -40 | -20 | Right temporal occipital fusiform cortex |
| | | 6.64 | 26 | -37 | -23 | Right temporal fusiform cortex pos |
| | | 6.51 | -36 | -44 | -16 | Left temporal occipital fusiform cortex |
| | | 6.45 | -39 | -34 | -28 | Left temporal fusiform cortex pos |
| | | 6.28 | -44 | -37 | -20 | Left inferior temporal gyrus post |
| | | 6.15 | 41 | -77 | -18 | Right lateral occipital cortex inf |
| | | 6.03 | -39 | -20 | -26 | Left temporal fusiform cortex pos |
| | 630 | 4.77 | -44 | 30 | 4 | Left inferior frontal gyrus p tri |
| | | 4.68 | -39 | 13 | 22 | Left inferior frontal gyrus p ope |
| | | 4.56 | -56 | 40 | 2 | Left frontal pole |
| | | 4.56 | -44 | 23 | 14 | Left inferior frontal gyrus p tri |
| | | 4.32 | -36 | 40 | -6 | Left frontal pole |
| | | 4.3 | -56 | 30 | -3 | Left inferior frontal gyrus p tri |
| | | 4.12 | -39 | 13 | 34 | Left middle frontal gyrus |
| | | 3.92 | -54 | 38 | 17 | Left frontal pole |
| | 286 | 4.59 | 14 | -97 | 27 | Right occipital pole |
| | | 4.51 | 21 | -94 | 20 | Right occipital pole |
| | | 4.35 | 11 | -90 | 22 | Right occipital pole |
| | | 4 | 9 | -92 | 32 | Right occipital pole |
| | | 3.95 | 31 | -100 | 12 | Right occipital pole |
| | 118 | 4.55 | -16 | -97 | 0 | Left occipital pole |
| | | 3.96 | -26 | -92 | 10 | Left occipital pole |
| | 413 | 4.42 | 44 | -40 | 42 | Right supramarginal gyrus pos |
| | | 4.28 | 54 | -37 | 42 | Right supramarginal gyrus pos |
| | | 4.23 | 49 | -37 | 32 | right superior longitudinal fas |

| | | | | | | |
|---|---|---|---|---|---|---|
| | | 3.98 | 44 | -42 | 52 | Right supramarginal gyrus pos |
| | | 3.86 | 29 | -47 | 42 | Right superior parietal lobule |
| | | 3.61 | 61 | -47 | 44 | Right angular gyrus |
| | | 3.5 | 59 | -37 | 34 | Right supramarginal gyrus pos |
| | 91 | 4.39 | -41 | 36 | -20 | Left frontal pole |
| | 93 | 4.17 | 64 | -24 | 20 | Right parietal operculum cortex |
| | | 3.99 | 66 | -27 | 27 | Right supramarginal gyrus ant |
| | 196 | 4.08 | -16 | -90 | 27 | Left occipital pole |
| | | 4.03 | -1 | -94 | 22 | Left occipital pole |
| | | 3.97 | -4 | -87 | 24 | Left cuneal cortex |
| | | 3.91 | -26 | -94 | 30 | Left occipital pole |
| | | 3.88 | -29 | -87 | 27 | Left lateral occipital cortex sup |
| | 85 | 3.94 | -1 | -50 | 52 | Left precuneous cortex |
| | | 3.7 | 11 | -60 | 54 | Right precuneous cortex |
| | | 3.26 | -4 | -62 | 57 | Left precuneous cortex |
| MEdn > ME | 8383 | 7.46 | 36 | -32 | -23 | Right temporal fusiform cortex pos |
| | | 7.22 | 26 | -37 | -23 | Right temporal fusiform cortex pos |
| | | 6.76 | 34 | -40 | -26 | Right temporal occipital fusiform cortex |
| | | 6.61 | 41 | -77 | -18 | Right lateral occipital cortex inf |
| | | 6.54 | -39 | -37 | -26 | Left temporal fusiform cortex pos |
| | | 6.53 | -36 | -44 | -16 | Left temporal occipital fusiform cortex |
| | | 6.51 | 21 | -47 | -18 | Right temporal occipital fusiform cortex |
| | | 6.45 | 29 | -52 | -16 | Right temporal occipital fusiform cortex |
| | 263 | 4.93 | 14 | -97 | 27 | Right occipital pole |
| | | 4.37 | 4 | -92 | 32 | Right occipital pole |
| | | 4.32 | -1 | -94 | 24 | Left occipital pole |
| | | 4.22 | 31 | -100 | 12 | Right occipital pole |
| | | 4.03 | -19 | -100 | 27 | Left occipital pole |
| | | 3.78 | 26 | -97 | 22 | Right occipital pole |
| | | 3.75 | -19 | -92 | 34 | Left occipital pole |
| | | 3.6 | -4 | -87 | 40 | Left cuneal cortex |

| | | | | | | |
|---|---|---|---|---|---|---|
| | 102 | 4.92 | -11 | 18 | 50 | Left superior frontal gyrus |
| | | 3.64 | -6 | 20 | 42 | Left paracingulate gyrus |
| | | 3.56 | -1 | 23 | 50 | Left superior frontal gyrus |
| | 129 | 4.74 | 39 | 23 | 14 | Right inferior frontal gyrus p tri |
| | | 3.71 | 59 | 30 | 10 | Right inferior frontal gyrus p tri |
| | | 3.67 | 56 | 28 | 20 | Right inferior frontal gyrus p tri |
| | 88 | 4.25 | 29 | -37 | 14 | Right lateral ventricle |
| | | 3.98 | 21 | -27 | 20 | Right lateral ventricle |
| | | 3.48 | 26 | -32 | 7 | Right lateral ventricle |
| | | 3.4 | 34 | -44 | 10 | right inferior frontal occipital fas |
| ME > MEdn | 80 | 4.54 | -71 | -20 | 2 | Left superior temporal gyurs pos |
| | | 3.76 | -61 | -24 | 7 | Left planum temporale |
| | | 3.23 | -71 | -30 | 7 | Left superior temporal gyurs pos |
| | 112 | 4.42 | 1 | -14 | 27 | Right cingulate gyrus ant |
| | | 3.73 | 9 | -24 | 34 | right cingulum cingulate |
| | | 3.69 | 6 | -22 | 24 | Right lateral ventricle |
| | 163 | 4.22 | -1 | 30 | 14 | Left cingulate gyrus ant |
| | | 4.09 | 1 | 38 | -6 | Right cingulate gyrus ant |
| | | 3.81 | -4 | 36 | 7 | Left cingulate gyrus ant |
| | | 3.65 | 6 | 46 | 0 | Right paracingulate gyrus |
| | | 3.36 | 16 | 48 | 2 | forceps minor |
| | 134 | 4.03 | 21 | -42 | -30 | Right corticospinal tract |
| | | 4.01 | 16 | -57 | -48 | Right cerebellum |
| | | 3.59 | 14 | -47 | -26 | Right cerebellum |
| | | 3.4 | 19 | -52 | -38 | Right corticospinal tract |
| 2 x 2 factorial design – echo and (downsampled) band | | | | | | |
| ANOVA | 486 | 5.4 | -44 | -40 | -18 | Left temporal fusiform cortex pos |
| | | 4.85 | -41 | -47 | -10 | left inferior longitudinal fas |
| | | 4.79 | -51 | -44 | -13 | left superior longitudinal fas |
| | | 4.58 | -46 | -52 | -20 | Left inferior temporal gyrus temocc |
| | | 4.44 | -34 | -44 | -20 | Left temporal fusiform cortex pos |
| | | 4.16 | -39 | -24 | -20 | Left temporal fusiform cortex pos |

| | | | | | |
|---|---|---|---|---|---|
| | 4.02 | -36 | -52 | -18 | Left temporal occipital fusiform cortex |
| | 3.82 | -59 | -40 | -10 | Left middle temporal gyrus pos |
| 281 | 5.39 | 44 | -34 | -23 | Right inferior temporal gyrus post |
| | 5.19 | 36 | -37 | -28 | Right temporal fusiform cortex pos |
| | 3.77 | 49 | -47 | -16 | Right inferior temporal gyrus temocc |
| | 3.74 | 39 | -24 | -18 | Right temporal fusiform cortex pos |
| | 3.7 | 34 | -40 | -18 | Right temporal fusiform cortex pos |
| | 3.28 | 36 | -54 | -20 | Right temporal occipital fusiform cortex |
| | 3.21 | 26 | -42 | -16 | Right temporal occipital fusiform cortex |
| 78 | 4.63 | -19 | 36 | -13 | Left frontal orbital cortex |
| | 3.5 | -19 | 28 | -18 | Left frontal orbital cortex |
| | 3.35 | -9 | 23 | -13 | Left subcallosal cortex |
| 57 | 4.39 | 31 | 36 | -10 | Right frontal pole |
| | 3.22 | 21 | 38 | -13 | Right frontal pole |
| 116 | 4.37 | -36 | 18 | 30 | Left middle frontal gyrus |
| | 4.13 | -36 | 13 | 17 | Left frontal operculum cortex |
| | 3.88 | -34 | 20 | 20 | Left inferior frontal gyrus p ope |
| | 3.83 | -46 | 26 | 24 | Left middle frontal gyrus |
| 64 | 4.19 | 41 | -70 | 10 | Right lateral occipital cortex inf |
| | 4.01 | 44 | -60 | 12 | Right lateral occipital cortex inf |

Supplementary table 2: Significant cluster and peak information for main effects, ANOVA effects of interest and directed effects of interest when comparing activation magnitude and activation precision. All coordinates are given in MNI space and all anatomical labels are extracted from the Harvard-Oxford cortical and subcortical structural atlases.

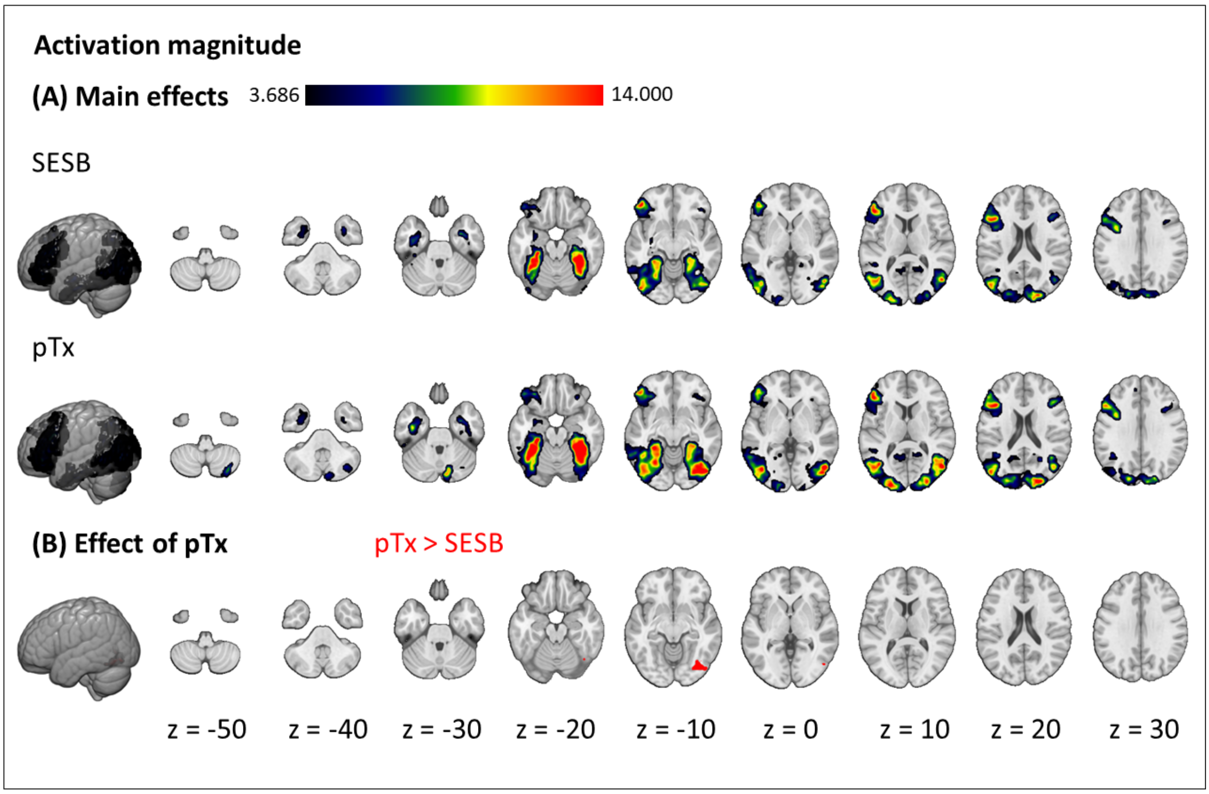


Supplementary figure 1: Effects of parallel transmit on activation magnitude. (A) effects of the SESB sequence and the pTx sequence for the contrast of interest ( S > C); (B) effect of parallel transmit (pTx > SESB, red; SESB > pTx, no significant clusters). Results are cluster-corrected at $p < 0.05$ based on an uncorrected voxel threshold of $p < 0.001$ and are overlaid on the MNI152NLin2009cAsym template.

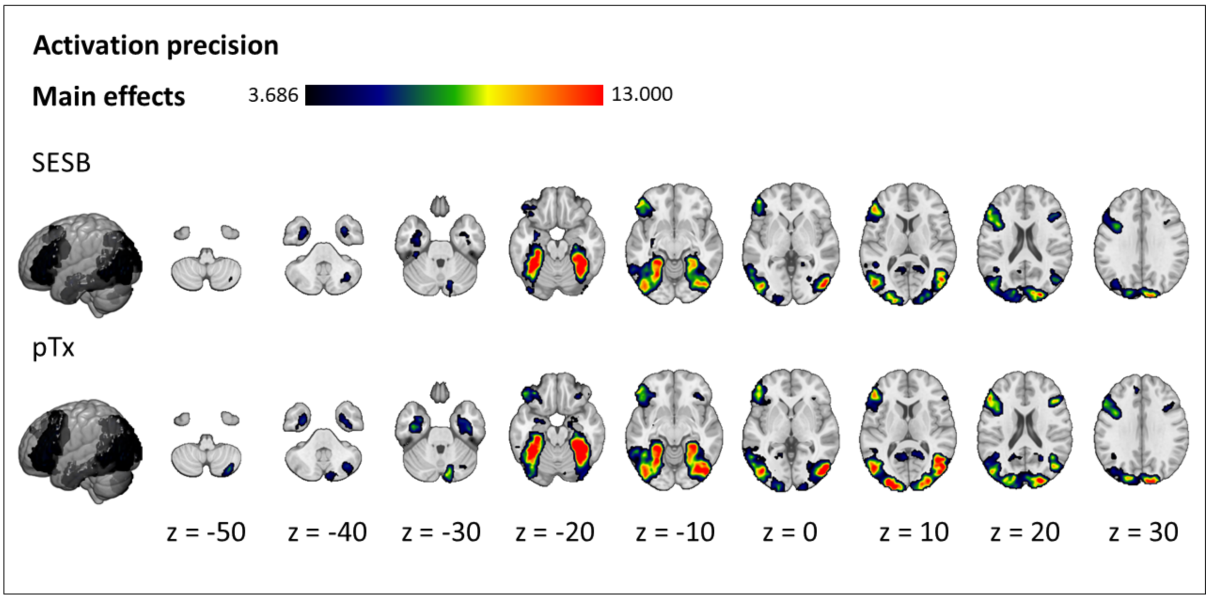


Supplementary figure 2: Effects of parallel transmit on activation precision. Effects of the SESB sequence and the pTx sequence for the contrast of interest (S > C). Results are cluster-corrected at $p < 0.05$ based on an uncorrected voxel threshold of $p < 0.001$ and are overlaid on the MNI152NLin2009cAsym template.

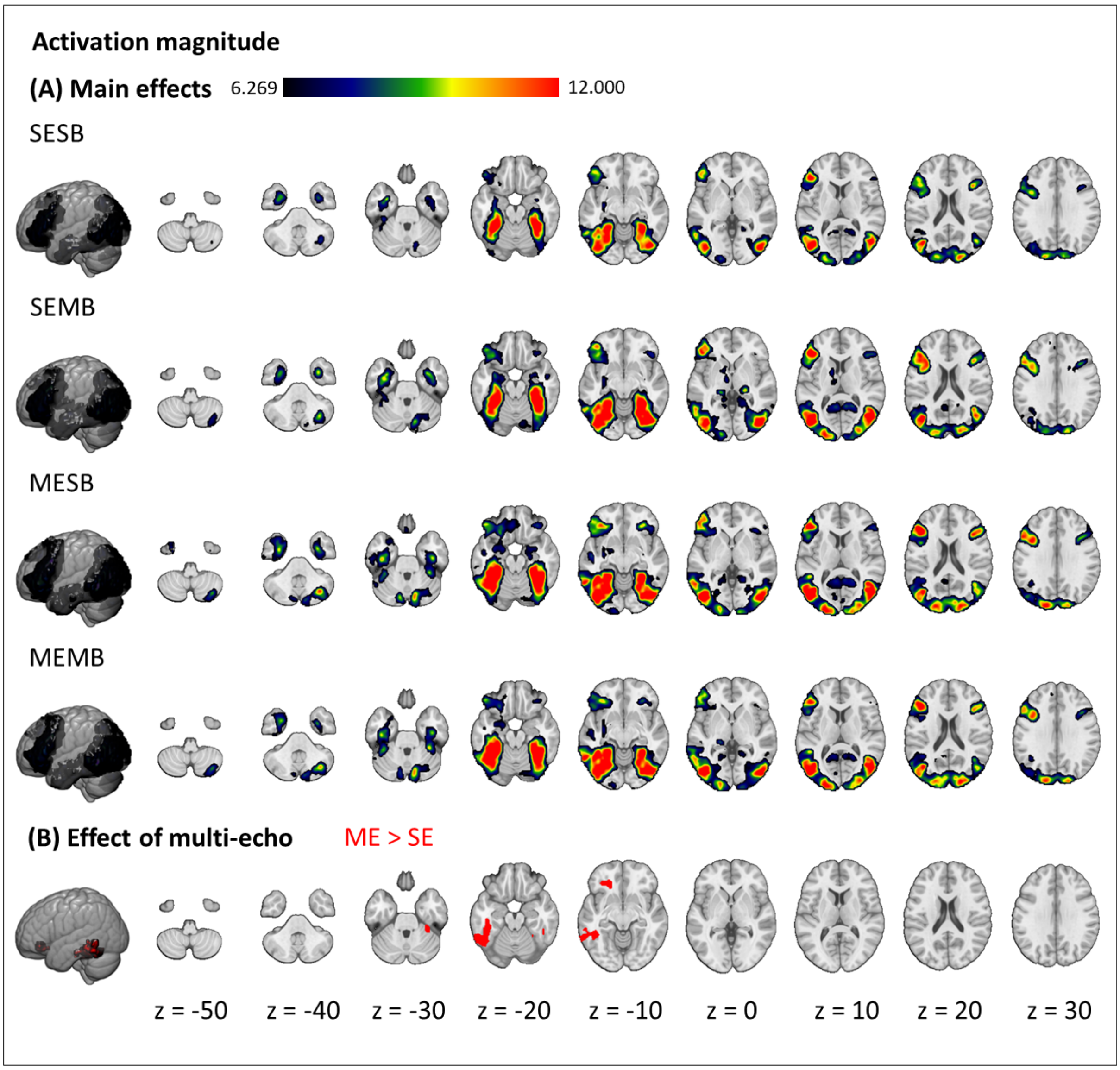


Supplementary figure 3: Effects of multi-echo and multiband on activation magnitude. (A) main effect of each sequence for the contrast of interest (S > C); (B) effect of echo (ME > SE, red; SE > ME, no significant clusters). Results are cluster-corrected at $p < 0.05$ based on an uncorrected voxel threshold of $p < 0.001$ and are overlaid on the MNI152NLin2009cAsym template.

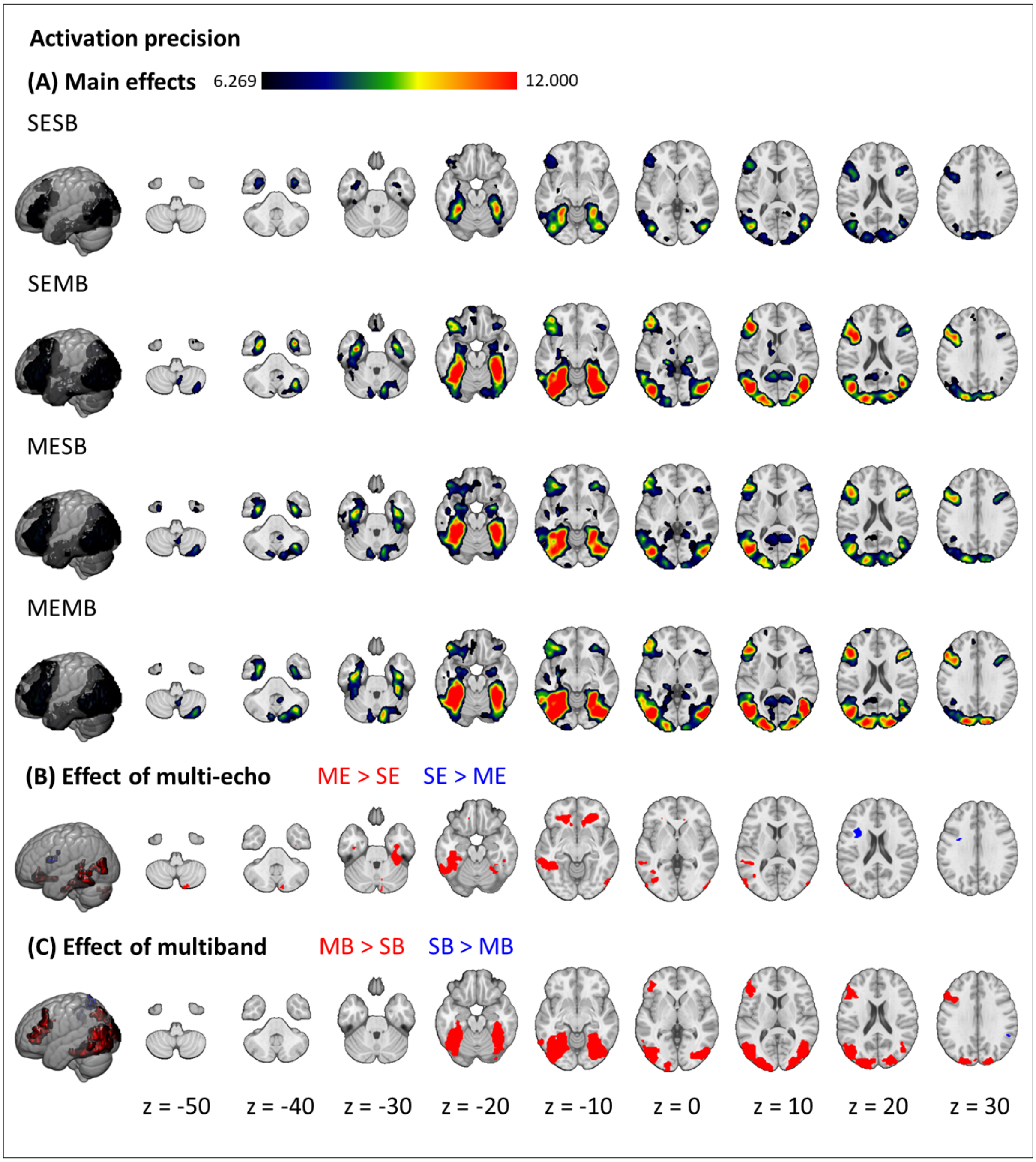


Supplementary figure 4: Effects of multi-echo and multiband on activation precision. (A) main effect of each sequence (S > C); (B) effect of echo (ME > SE, red; SE > ME, blue); (C) effect of band (MB > SB, red; SB > MB, blue). Results are cluster-corrected at $p < 0.05$ based on an uncorrected voxel threshold of $p < 0.001$ and are overlaid on the MNI152NLin2009cAsym template.

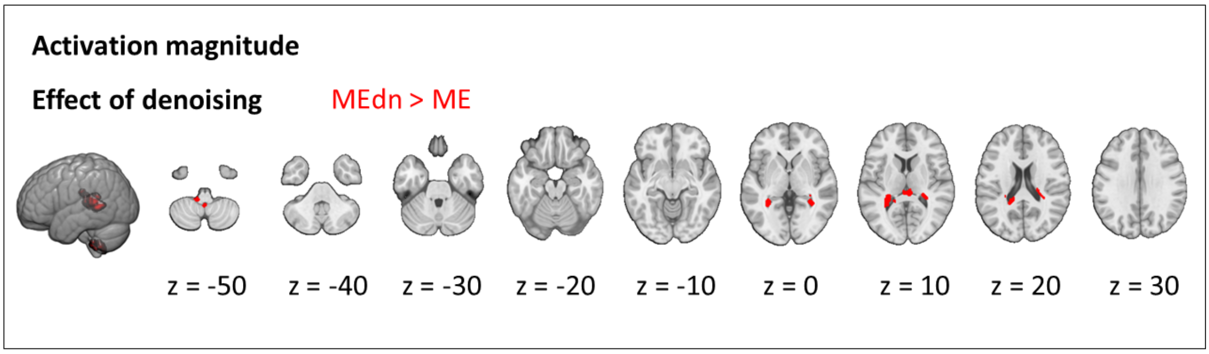


Supplementary figure 5: Effect of ME-ICA denoising on activation magnitude (MEdn > ME, red; ME > MEdn, no significant clusters). Results are cluster-corrected at $p < 0.05$ based on an uncorrected voxel threshold of $p < 0.001$ and are overlaid on the MNI152NLin2009cAsym template.

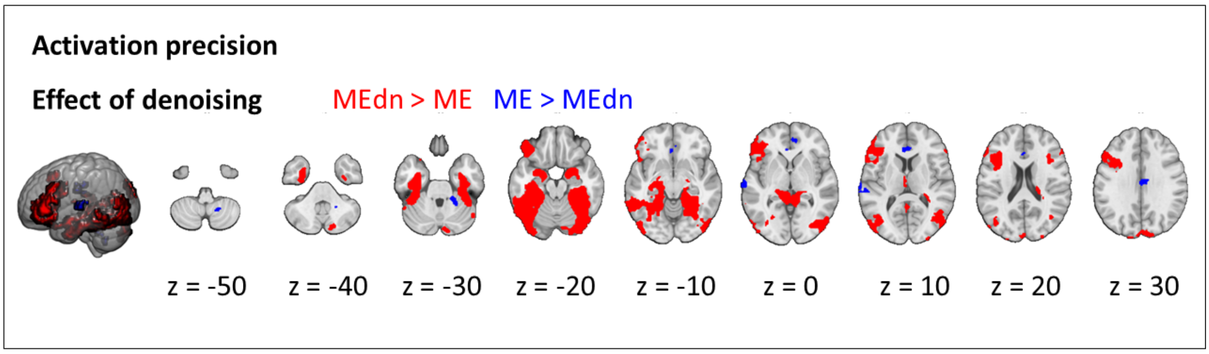


Supplementary figure 6: Effect of ME-ICA denoising on activation precision (MEdn > ME, red; ME > MEdn, blue). Results are cluster-corrected at $p < 0.05$ based on an uncorrected voxel threshold of $p < 0.001$ and are overlaid on the MNI152NLin2009cAsym template.

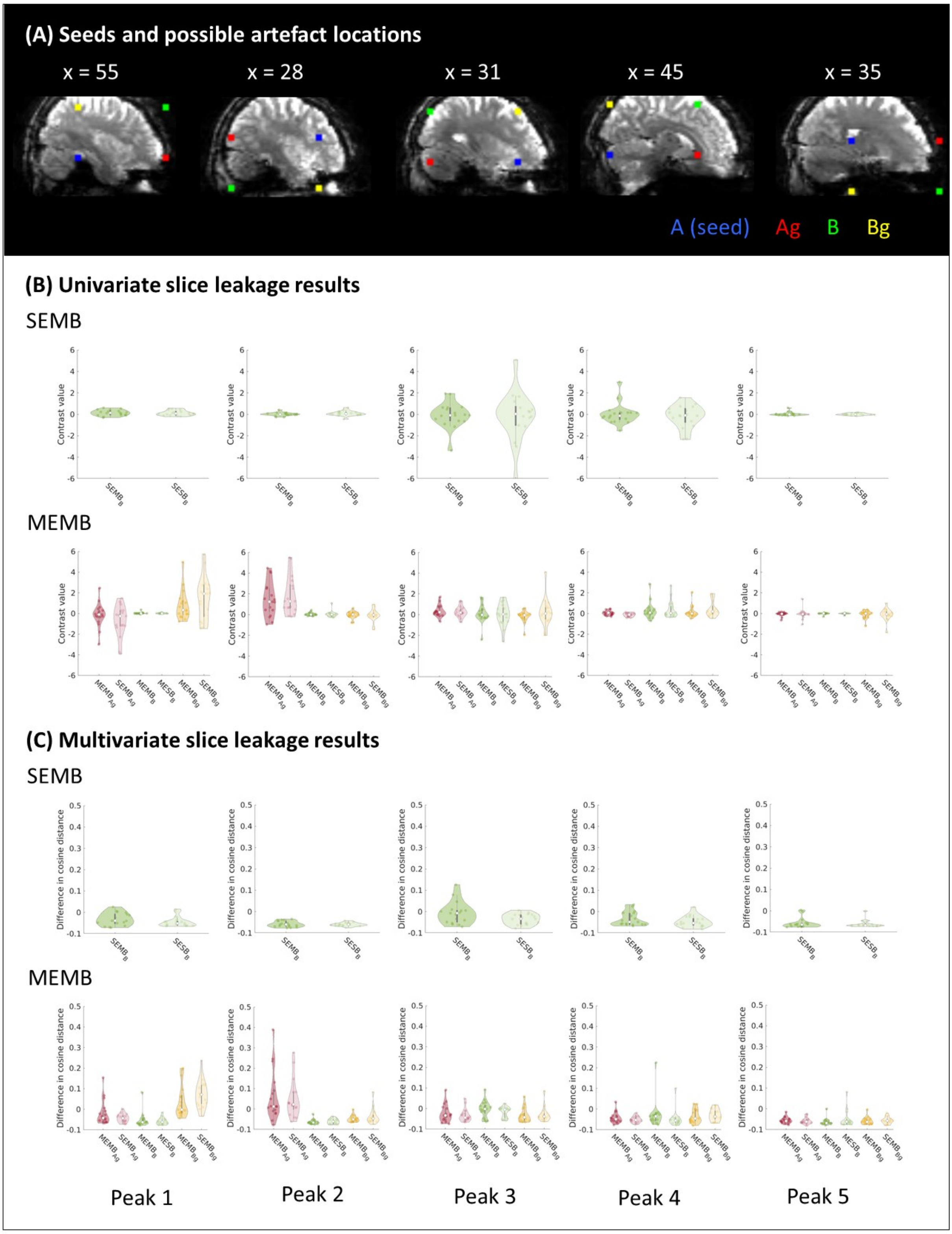


Supplementary figure 7: Slice leakage analysis (all peaks). (A) Seed and possible artefact locations for a single participant. The ROIs (radius 4 voxels) indicate the seed location (blue), possible artefact location based on phase shift (green), possible artefact location based on GRAPPA (red), and possible artefact location based on phase shift and GRAPPA (yellow) in native EPI space. Informed consent was obtained from the participant for this image to be published. (B) Mean activation magnitude within each sphere for each MB sequence and the corresponding control sequence for each seed location. (C) Mean MVPA dissimilarity within each sphere for each MB sequence and the corresponding control sequence.

| | Peak 1 | Peak 2 | Peak 3 | Peak 4 | Peak 5 |
|---|---|---|---|---|---|
| Activation magnitude | | | | | |
| $SEMB_B$ | 0.3228 | 0.8153 | 0.3865 | 0.2161 | 0.6244 |
| $MEMB_{Ag}$ | 0.0539 | 0.9614 | 0.3099 | 0.0068* | 0.5378 |
| $MEMB_B$ | 0.2499 | 0.5327 | 0.3019 | 0.7382 | 0.5362 |
| $MEMB_{Bg}$ | 0.9647 | 0.3988 | 0.8770 | 0.9917 | 0.5296 |
| MVPA | | | | | |
| $SEMB_B$ | 0.0596 | 0.3208 | 0.0107* | 0.1079 | 0.3597 |
| $MEMB_{Ag}$ | 0.0974 | 0.1860 | 0.2596 | 0.0959 | 0.3434 |
| $MEMB_B$ | 0.2528 | 0.6570 | 0.0354* | 0.0083* | 0.8442 |
| $MEMB_{Bg}$ | 0.9408 | 0.8805 | 0.5601 | 0.8335 | 0.5945 |

Supplementary table 3: p-values for all slice leakage tests. A is the seed location, B is the possible artefact location based on phase shift, Ag is the possible artefact location in the same slice as A based on GRAPPA, and Bg is the possible artefact location in the same slice as B based on GRAPPA. * = $p < 0.05$, ** = $p < 0.05$ (Bonferroni-corrected for the number of peaks and possible artefact regions), SEMB = single-echo multiband, multiband, MEMB = multi-echo multiband.
