## Supplementary Material 2 for "Optimising 7T-fMRI for imaging regions of magnetic susceptibility"

**Supplementary Material 2: Investigating the impact of Ernst angle on the effects of interest**

### 1. Introduction

Due to experimenter error, the flip angle for our SESB and SEMB was not set to the Ernst angle (SESB: nominal flip angle = 50° compared to Ernst angle of 78°; SEMB: nominal flip angle of 50° compared to Ernst angle of 63°). Theoretically, use of suboptimal flip angles would result in a tSNR loss relative to using the Ernst angle – 12.99% for the SESB sequence and 3.15% for the SEMB sequence – and this loss might explain the poorer performance of SE sequences relative to ME sequences. However, because of $B_1$ inhomogeneity at 7T, the effective flip angle may differ from the nominal flip angle and so the actual signal loss may differ from theoretical predictions. This difference may vary across the brain, especially for CP-mode scans (Uğurbil, 2018), so it is important to determine the actual effect within ventral temporal and orbitofrontal regions. We acquired data to test the impact of suboptimal flip angle on tSNR. Since tSNR is known to dissociate from functional contrast under some circumstances (Ding et al., 2022), we also tested the impact on our effects of interest (activation magnitude, activation precision, and our MVPA metric) within our ROIs.

### 2. Methods

We collected data from two healthy participants who did not participate in the main study and who gave informed consent. There were four functional runs per participant, collected in a single session. The first two runs used the SE sequences described in the main text (Run 1 – SESB, Run 2 – SEMB). The other two runs used SE sequences in which the nominal flip angle was set to the Ernst angle (all other parameters were kept constant between matched sequences; Run 3 – SESBernst, Run 4 - SEMBernst). All other aspects of the methods, including task, preprocessing, and GLMs, were identical to those described in the main text. tSNR, activation magnitude, activation precision, and MVPA performance were calculated within every ROI reported in the main text.

### 3. Results

Supplementary table 4 shows the results. In both participants, use of the Ernst angle conferred a tSNR increase that was relatively consistent across ROIs. The difference between the suboptimal flip angle and the Ernst angle was greater for the SESB sequence than for the SEMB sequence; consistent with this, the improvement in tSNR when using the optimised sequence was greater for the SESB sequence. These tSNR results were therefore in line with what would be expected from a sequence using the Ernst angle. However, our three effects of interest – activation magnitude, activation precision, and our MVPA metric – showed an inconsistent pattern. The magnitude and direction of change when using the optimised sequence relative to the suboptimal sequence varied across ROIs and across participants.

| | LTP | LvATL | RITG | LFP | LmMTG | LpMTG | LIFGpt |
|---|---|---|---|---|---|---|---|
| **tSNR** | | | | | | | |
| Participant 1 | | | | | | | |
| SESBernst > SESB | 54.0286 | -9.9851 | 59.8135 | 19.7037 | 14.1672 | 4.3193 | 19.8550 |
| SEMBernst > SEMB | 34.4127 | -20.3186 | 3.6741 | 12.7745 | 3.0920 | 3.2001 | 13.1559 |
| Participant 2 | | | | | | | |
| SESBernst > SESB | -3.2123 | 19.6513 | 87.5284 | 52.9209 | 20.7838 | 24.4694 | -5.9168 |
| SEMBernst > SEMB | 24.7779 | -0.7088 | -7.1845 | 31.0988 | 0.3452 | 9.3827 | 1.0219 |
| **Activation magnitude** | | | | | | | |
| Participant 1 | | | | | | | |
| SESBernst > SESB | 44.0782 | 583.4712 | 227.3575 | -742.3157 | -211.2123 | 46.1007 | 1564.3744 |
| SEMBernst > SEMB | -3.2425 | -45.0988 | -85.7035 | -426.4440 | -36.8633 | -91.4717 | -99.7580 |
| Participant 2 | | | | | | | |
| SESBernst > SESB | -57.6951 | 11.2470 | -1.9989 | -75.8461 | -147.4325 | -55.1979 | -102.6642 |
| SEMBernst > SEMB | -36.7422 | -39.1730 | -16.8902 | 191.7208 | 82.8526 | -57.2460 | -7.2006 |
| **Activation precision** | | | | | | | |
| Participant 1 | | | | | | | |
| SESBernst > SESB | 63.3282 | 588.9795 | 219.5805 | -1345.4358 | -220.6662 | 73.0525 | 1588.8752 |
| SEMBernst > SEMB | -0.4183 | -37.5597 | -88.0863 | -412.9069 | -20.8991 | -88.6187 | -100.0000 |
| Participant 2 | | | | | | | |
| SESBernst > SESB | -65.4802 | 19.5963 | -7.2510 | -75.8659 | -155.2239 | -60.3942 | -101.5743 |
| SEMBernst > SEMB | -34.6221 | -41.4298 | 16.2403 | 133.2690 | 83.1263 | -52.9855 | -9.2875 |
| **MVPA** | | | | | | | |
| Participant 1 | | | | | | | |

| | | | | | | | |
|---|---|---|---|---|---|---|---|
| SESBernst > SESB | -23.9196 | 122.1070 | 47.6859 | 154.8006 | -32.3599 | 35.5384 | 35.1941 |
| SEMBernst > SEMB | 63.3551 | -111.4677 | -149.0596 | 50.2805 | -1.8231 | -297.4278 | -120.5662 |
| Participant 2 | | | | | | | |
| SESBernst > SESB | -107.5576 | -24.3165 | -109.6968 | -85.2973 | 282.0658 | -129.8579 | -128.2633 |
| SEMBernst > SEMB | -211.9854 | -89.6384 | 49.8018 | -19172.9623 | 5.6581 | -2058.3843 | -78.0659 |

Supplementary table 4: Impact of suboptimal flip angles on tSNR and effects of interest. Percentage change is percentage improvement using optimised sequence relative to sequence reported in main text. SESB = single-echo single band sequence reported in main text, SESBernst = single-echo single band sequence using Ernst angle as nominal flip angle, SEMB = single-echo multiband sequence reported in main text, SEMBernst = single-echo multiband sequence using Ernst angle as nominal flip angle, LTP = left temporal pole, LvATL = left ventral anterior temporal lobe, RITG = right inferior temporal gyrus, LFP = left frontal pole, LmMTG = left medial middle temporal gyrus, LpMTG = left posterior middle temporal gyrus, LIFGpt = left inferior temporal gyrus pars triangularis.

### 4. Discussion

To thoroughly eliminate all possible confounds, future studies investigating the effect of multi-echo sequences relative to single-echo sequences should strive to set the flip angle to the Ernst angle and to ensure that this is consistent across sequences. However, these analyses suggest that suboptimal flip angle alone cannot explain the benefits of ME over SE for our outcome effects of interest (activation magnitude, activation precision, and MVPA performance) that was observed in the main study. It is possible that a group difference would emerge in a well-powered study designed specifically to investigate the effect of flip angle; however, these preliminary results are sufficient to highlight large interindividual variability. These results also align with previous work that finds a dissociation between tSNR and functional contrast metrics (Ding et al., 2022).

We therefore conclude that the difference in flip angle is unlikely to explain the reported advantage of ME over SE sequences.

**Background**

7T-fMRI can offer researchers...

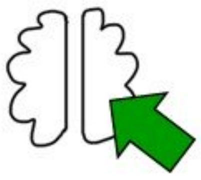

... better spatial specificity

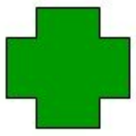

... shorter scan times for patients

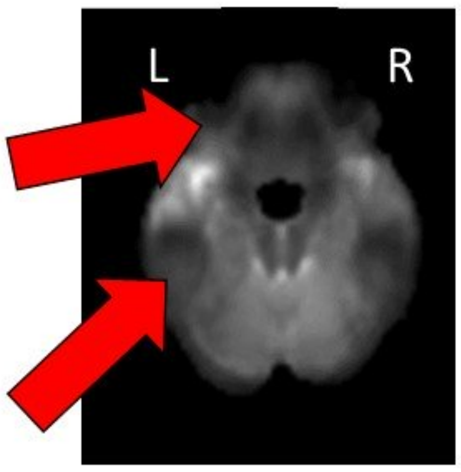


However, 7T is associated with exacerbated signal dropout and distortions in ventral temporal and orbitofrontal regions.

**Methods**

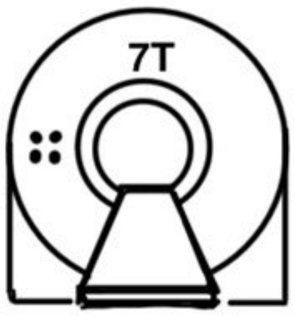


n = 20 healthy participants

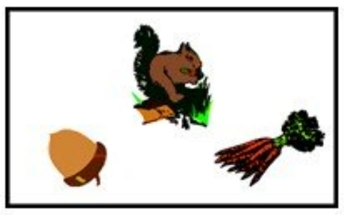

Semantic judgement task

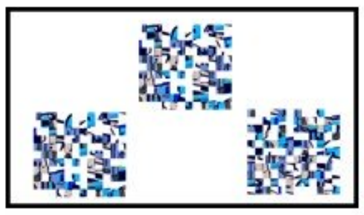

Pattern matching (control) task

The contrast of interest (semantic > control) includes temporal and orbitofrontal areas.

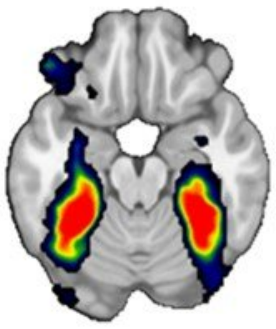

**Three methods for improving image quality:**

1. Parallel transmit (pTx)
2. Multi-echo (ME)
3. Multiband (MB)

**Two effects of interest:**

1. Activation magnitude (contrast betas)
2. Precision of GLM fit (contrast t-values)

**Results**

**Activation magnitude**

pTx > baseline

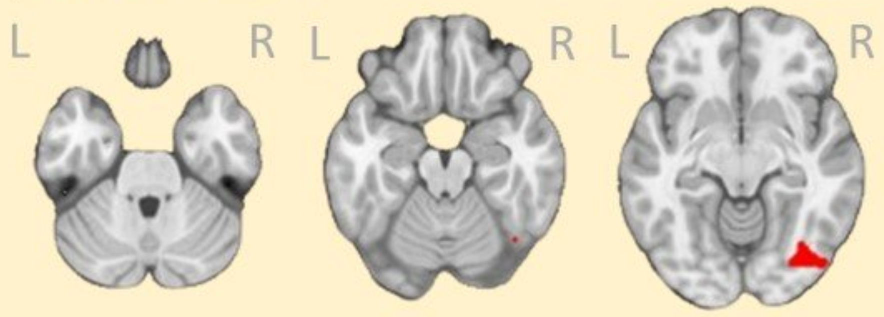


Multi-echo > single echo

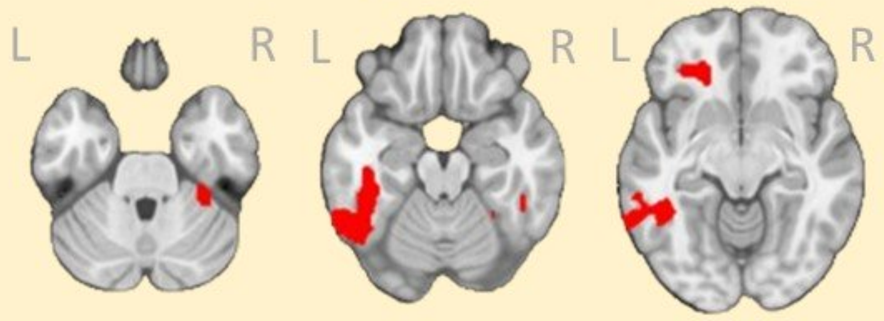


**Activation precision**

Multiband > single band

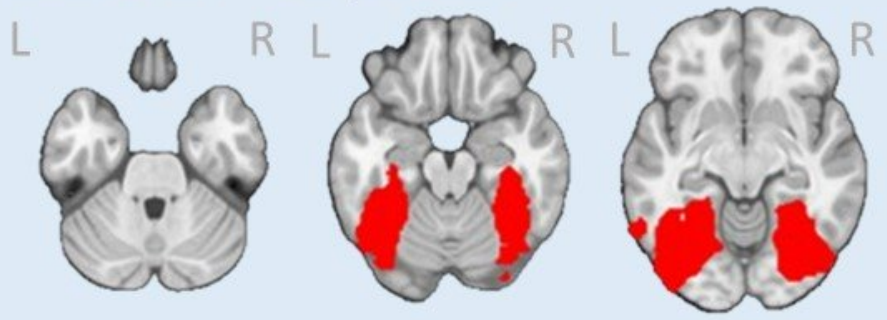


Multi-echo with ICA (tedana) > multi-echo

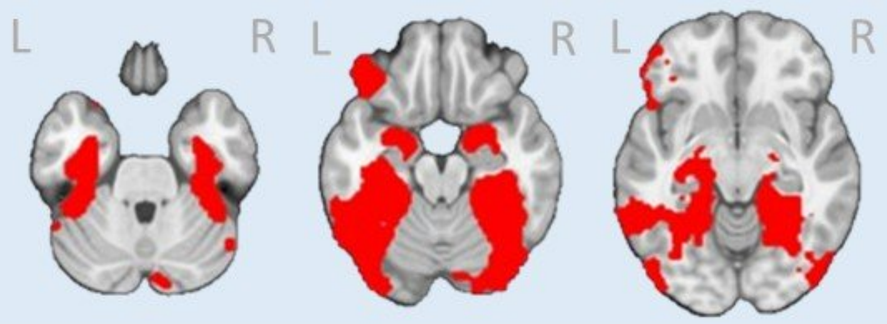


**Conclusion:** a multi-echo, multiband acquisition sequence was best at detecting signal in artefact-prone regions while maintaining image quality across the rest of the brain.
